# Novel piece of the puzzle: ALI1 is required for oxo-C14-HSL priming in Arabidopsis

**DOI:** 10.1101/2021.12.09.471921

**Authors:** Abhishek Shrestha, Casandra Hernández-Reyes, Maja Grimm, Johannes Krumwiede, Elke Stein, Sebastian T. Schenk, Adam Schikora

**Author notes:** Corresponding author: Adam Schikora.

## Abstract

Quorum sensing (QS) molecules mediate communication between bacterial cells. *N*-acyl homoserine lactones (AHL) are one of the best-studied groups of QS molecules. In addition to bacterial communication, AHL are involved in interactions with eukaryotes. Short side-chain AHL are readily taken up by plants. They induce root elongation and growth promotion. Hydrophobic long side-chain AHL are usually not transported over long distances although, they may prime plants for enhanced resistance. Unfortunately, studies elucidating the plant factors required for response to AHL are sparse. Here, we provide evidence of a plant protein, namely the <u>A</u>H<u>L</u>-priming protein 1 (ALI1), indispensable for enhanced resistance response induced by the *N*-3-oxotetradecanoyl-homoserine lactone (oxo-C14-HSL). Comparing Col-0 and the *ali1* mutant, we revealed loss of AHL-priming in *ali1*. This phenomenon is reverted with the reintroduction of *ALI1* into *ali1*. Additional transcriptome analysis revealed that *ali1* is less sensitive to oxo-C14-HSL treatment compared to the wild-type. Our results suggest, therefore, that ALI1 is required for oxo-C14-HSL-dependent priming for enhanced resistance in Arabidopsis.

## Introduction

Being in constant vicinity of potentially harmful microorganisms, plants depend on efficient perception mechanisms and swift responses. Unlike mammals, which possess a repertoire of mobile defense cells, immune responses in plants rely on their individual cells and their potential to recognize microorganisms and respond to them. The first level of recognition is mediated through membrane receptor proteins called pattern recognition receptors (PRRs). They perceive microbe-associated molecular patterns (MAMPs), which are conserved microbial structural molecules like flagellin, glycoprotein or chitin (Gomez-Gomez and Boller, 2000; Zipfel et al., 2006; Miya et al., 2007; Gohre and Robatzek, 2008). The recognition of MAMPs by PRRs leads to the activation of innate immune response termed as pattern-triggered immunity (PTI). PTI is associated with activation of mitogen-activated protein kinase (MAPK), transcriptional induction of defense-related genes, production of reactive oxygen species (ROS) and callose deposition at sites of infection; all inhibit microbial invasion and growth (De Wit, 1997; Nürnberger et al., 2004; Schenk et al., 2014).

Another nature of microbial interaction with higher organisms is mediated through quorum sensing (QS) molecules, such as *N*-acyl homoserine lactones (AHL). Gram-negative bacteria generally rely on the synthesis of autoinducers like AHL or cyclodipeptides to orchestrate their cell behavior by activating and impeding transcription of QS-regulated genes (Miller and Bassler, 2001; Geske et al., 2008; Ni et al., 2009). Besides orchestrating the communication within bacterial populations, quorum sensing (QS) molecules like AHL modulate interactions between bacteria and eukaryotes. AHL comprise of two intrinsic moieties, a core *N*-acylated homoserine lactone ring and an amide linked acyl side-chain that varies in length from 4 to 18 carbons and in the substitution of the hydrogen with a hydroxyl or ketone group on the third carbon position (Whitehead et al., 2001; Marketon et al., 2002; von Bodman et al., 2003).

Since eukaryotes coevolved with bacteria, it is maybe not surprising that QS molecules elicit specific responses (Williams, 2007). Numerous studies have assessed this phenomenon in mammalian cells, primarily focused on the effects of *N*-3-oxododecanoyl-*L*-homoserine lactone (oxo-C12-HSL), an AHL produced by pathogenic human pathogen *Pseudomonas aeruginosa*. Kravchenko et al. (2006) showed that the perception of AHL in animal cells is independent of canonical innate immune system receptors. Hydrophobic oxo-C12-HSL freely diffuses the membranes of T-cells and model lipid membranes (Ritchie et al., 2007; Barth et al., 2012) through membrane microdomains such as caveolae and lipid rafts (Davis et al., 2010). AHL activates diverse signaling components and pathways in mammalian cells, including calcium signaling, activation of Rho GTPases, MAPK and transcription factor NFκB, cytokines, chemokines, enzymes and interferons (Smith et al., 2002; Shiner et al., 2006; Kravchenko et al., 2008; Mayer et al., 2011; Karlsson et al., 2012a; Glucksam-Galnoy et al., 2013). Peroxisome proliferator-activated receptors PPARγ and PPARβ, members of the nuclear hormone receptor (NHR) family, were proposed as potential candidates for AHL-interacting proteins in mammalian cells (Jahoor et al., 2008), although a confirmation of physical binding was missing. Contrarily, oxo-C12-HSL was shown to interact and colocalize with the IQ-motif containing GTPase-activating protein1 (IQGAP1) that leads to changes in phosphorylation of Rac1 and Cdc42 (Karlsson et al., 2012b). Moreover, oxo-C12-HSL from *P. aeruginosa* displayed association and activation of the bitter receptor T2R38 on leucocytes of different origins (Maurer et al., 2015; Gaida et al., 2016).

Plants respond to the presence of short-chained AHL like *N*-hexanoyl-*L*-homoserine lactone (C6-HSL) and *N*-octanoyl-*L*-homoserine lactone (C8-HSL) with growth modifications, including changes in root morphology and enhanced plant biomass in Arabidopsis (von Rad et al., 2008; Schikora et al., 2011; Schenk et al., 2012; Shrestha et al., 2020) and crop plants like wheat and barley (Rankl et al., 2016; Moshynets et al., 2019). These growth modulations were shown to depend on auxin signaling processes (von Rad et al., 2008; Bai et al., 2011; Liu et al., 2012). Another AHL, *N*-3-oxodecanoyl-*L*-homoserine lactone (oxo-C10-HSL), triggered the formation of adventitious roots in mung beans by inducing shortening of roots but augmented lateral root and root hair formations. This effect was induced through enhanced basipetal auxin transport followed by the accumulation of hydrogen peroxide and nitric oxide (Bai et al., 2011). However, some studies showed AHL-mediated modulation of plant development without the involvement of auxin (Ortiz-Castro et al., 2008; Palmer et al., 2014). The long-chained AHL induce so-called AHL-priming for enhanced resistance in which primed plants elicited faster and stronger immune responses than naive plants (Zarkani et al., 2013; Hernández-Reyes et al., 2014; Shrestha et al., 2019). Furthermore, *Arabidopsis thaliana* and barley (*Hordeum vulgare*) when inoculated with the *N*-3-oxotetradecanoyl-*L*-homoserine lactone (oxo-C14-HSL)-producing *Ensifer meliloti* (*Sinorhizobium meliloti*) strain *expR*+ displayed enhanced resistance to *Pseudomonas syringae* and *Blumeria graminis,* respectively compared to control plants (Zarkani et al., 2013; Hernández-Reyes et al., 2014; Shrestha et al., 2019). So far, different defense hormones are shown to be involved in AHL-induced systemic resistance, including salicylic acid and oxylipin 12-oxophytodienoic acid (cis-OPDA) (Hartmann et al., 2004; Schuhegger et al., 2006; Schikora et al., 2011; Schenk et al., 2014). Oxo-C14-HSL induced priming for enhanced defense was also associated with augmented and prolonged activation of the MAP kinases, MPK3 and MPK6, along with the upregulation of defense-related genes *WRKY22*, *WRKY29*, *GST6* and *Hsp70* (Beckers et al., 2009; Schikora et al., 2011; Schenk and Schikora, 2015; Shrestha et al., 2019). Furthermore, in response to triggering stress, primed plants exhibit an increased accumulation of reactive oxygen species (ROS), phenolic compounds and callose in their cell walls (Schenk and Schikora, 2015).

Several studies elucidated components that mediate AHL response in plants. Liu et al. (2012) discovered that G-protein coupled receptor GCR1 and the canonical Gα subunit, GPA1, are required for *N*-3-oxohexanoyl-*L*-homoserine lactone (oxo-C6-HSL) and *N*-3-oxooctanoyl-*L*-homoserine lactone (oxo-C8-HSL) mediated root elongation effects in *Arabidopsis*. Additionally, *N*-butanoyl-*L*-homoserine lactone (C4-HSL) was shown to induce Ca^2+^ spiking in *Arabidopsis* root cells (Song et al., 2011). The same group postulated the involvement of calmodulin proteins in response to C6-HSL in *Arabidopsis* (Zhao et al., 2015). AHL-mediated growth-promoting effect was shown to be dependent on the generation of a byproduct, *L*-homoserine, after AHL amidolysis, which increases the transfer of minerals and water through enhanced transpiration (Palmer et al., 2014). Different studies have demonstrated that the short-chain AHL molecules are readily taken up through roots and transported to shoots in plants, while the long-chain AHL molecules are not due to their lipophilic nature (Götz et al., 2007; von Rad et al., 2008; Sieper et al., 2014).

In the present study, we investigated one of such components, a protein critical for oxo-C14-HSL-mediated enhanced resistance in plants. We propose the Arabidopsis protein AHL-Priming Protein 1 (ALI1), previously annotated as ATGALK2 (GALACTOKINASE 2) to be an integral component for oxo-C14-HSL-mediated priming in Arabidopsis.

## Results

### Principal mechanism of AHL-induced priming is missing in *ali1*

To identify plant factors required for efficient AHL-priming, we focused on the gene encoded by the *At5g14470* locus since this gene was identified in previous AHL-priming-related studies. *At5g14470* locus encodes a putative kinase from the GHMP kinase family, previously named ATGALK2 or GALACTOKINASE 2. However, because of unclarities in the current annotation and the function prediction, we propose to rename it to <u>A</u>H<u>L</u>-Priming Protein <u>1</u> (ALI1).

To verify if oxo-C14-HSL influences the expression of *ATGALK2*/*ALI1*, quantitative PCR analysis was carried out with wild-type Col-0 seedlings that were pretreated with 6 μM oxo-C14-HSL for three days and subsequently elicited with 100 nM flg22. The results revealed that pretreatment with oxo-C14-HSL had no impact on the expression of *At5g14470* locus encoded gene, when compared to the control (acetone) treated Col*-*0 plants. However, 2 h after flg22 elicitation, we could observe two folds increase in *ATGALK2/ALI1* expression level in the Col*-*0 plants, irrespective of the pretreatment. Furthermore, the enhanced expression of *ATGALK2/ALI1* in Col*-* 0 due to flg22 challenge appears to be transient since the expression level returned to the basal level 24 h after flg22 challenge, irrespective of the pretreatment (Supplemental Fig. S1). This suggests that ATGALK2/ALI1 might play a role in the early response to bacteria.

To further corroborate on the role of ATGALK2/ALI1 during AHL-priming, we used a mutant line with T-DNA insertion in *At5g14470* locus, along with wild-type Col-0. The segregating line N560407 was obtained from Salk Institute, propagated and a non-segregating line (*ali1*) was characterized in a PCR approach (Supplemental Fig. S2). We performed quantitative PCR and western blot-based kinase assays to verify whether AHL-priming depends on the presence of functional ATGALK2/ALI1 in Arabidopsis (Schikora et al., 2011). Higher expression of *TLP5* and *DFR* genes, usually observed upon AHL-priming (Schenk *et al*, 2014) was missing in the *ali1* mutant (Fig. 1A). Furthermore both, Col-0 and the *ali1* mutant responded to the flg22 challenge and activated MPK6 transiently. In control Col-0 plants, the phosphorylation on the MPK6 activation loop was detectable 30 min after challenge with 100 nM flg22 but the phosphorylation level declined 60 min post challenge. No phosphorylation of the activation loop was observed 120 min after the flg22 challenge. On the contrary, oxo-C14-HSL-pretreated plants exhibited enhanced and prolonged activation of MPK6 since the pTEpY epitope was observed until 120 min after the flg22 challenge (Fig. 1B). In *ali1* mutant pretreated with either acetone or oxo-C14-HSL, the pattern of MPK6 phosphorylation was identical, and similar to that of control Col-0 plants (Fig. 1B). These results indicate that the ATGALK2/ALI1 is required for the oxo-C14-HSL-mediated enhanced and prolonged activation of MPK6 in response to flg22 challenge in AHL-primed plants.

**Figure 1.**
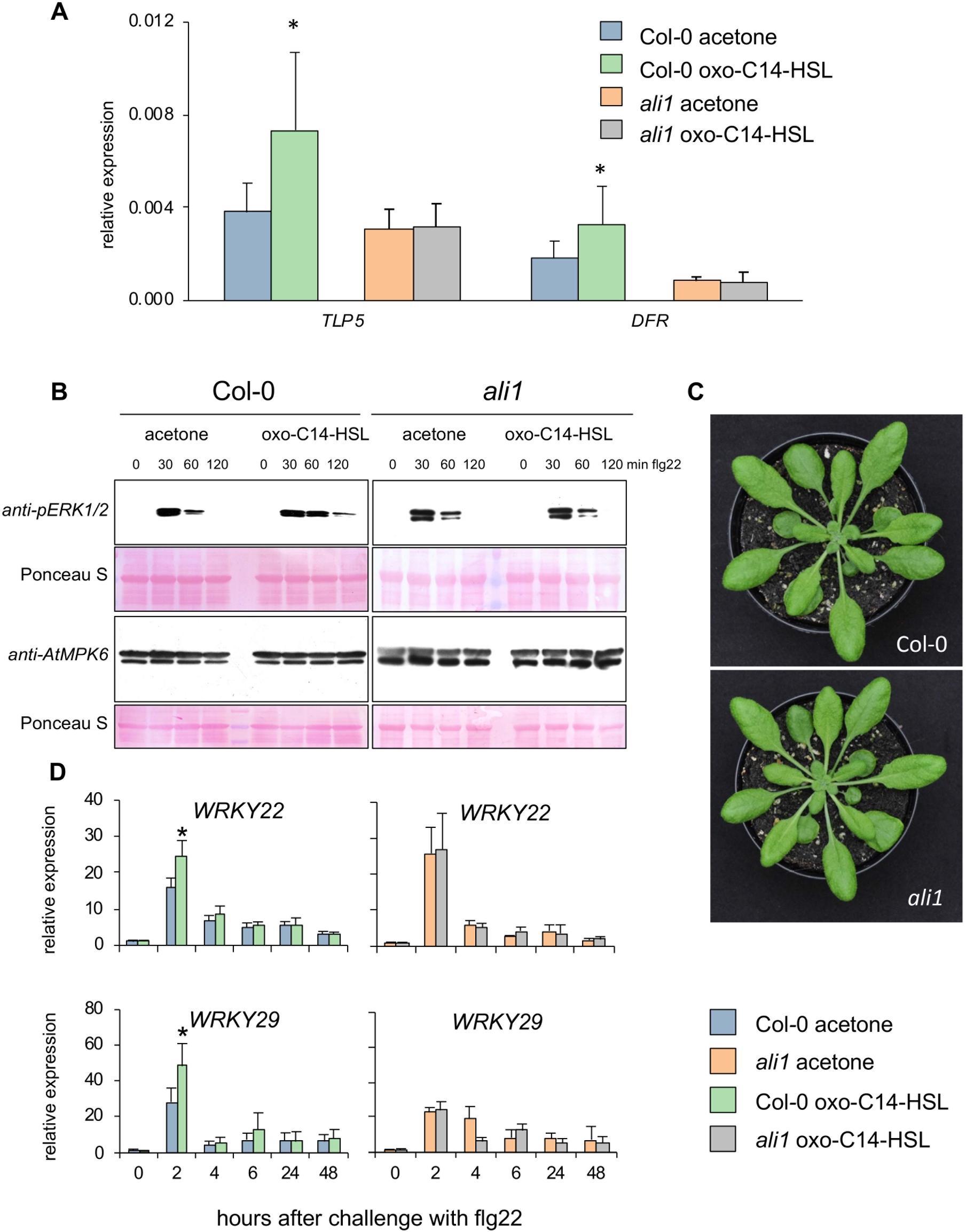
Principal mechanism of AHL-induced priming is missing in *ali1.* A, Expression analysis of two AHL-priming marker genes (*TLP5* and *DLR*) in Col-0 and *ali1* after priming with oxo-C14-HSL. The abundance of each gene transcript was normalized with Ubiquitin ligase (*At5g25760*) transcript and 0 hpt (hours post treatment) levels. The bar represents mean and error bars represent SD from eight independent biological repetitions from three experimental replicates. B, Phosphorylation pattern of the AtMPK6 in response to flg22 challenge in Col-0 and *ali1*, pretreated with 6 μM oxo-C14-HSL or acetone (control). Western blot analysis was performed in three independent experiments, representative blots are shown. C, Representative images of Col-0 wild type and the *ali1* mutant. D, Expression profile of defense-related genes *WRKY22* and *WRKY29* was monitored at various time points after 100 nM flg22 challenge in Col-0 and *ali1* mutant. The abundance of each gene transcript was normalized with *Ubiquitin ligase* (*At5g25760*) transcript and 0 hpt (hours post treatment) levels. Error bars represent SD from four independent biological repetitions. Experiments were carried out three times with similar results. Data information: In (A and D), data are presented as mean ± SD. * *P* ≤ 0.05 (Student’s *t*-test).

To determine whether AHL-priming reflected by enhanced upregulation of defense-related genes after a challenge (Schikora et al., 2011; Schenk and Schikora, 2014; Shrestha et al., 2019) is influenced by the absence of *ALI1*, we assessed genes’ expression using a qPCR approach. Analysis of *WRKY22* and *WRKY29* expression in Col-0 and *ali1* revealed that the seedlings were able to perceive 100 nM flg22 and elicit a response (Fig. 1D). However, the comparison between Col-0 and *ali1* revealed higher transcript abundance in oxo-C14-HSL-pretreated Col-0 seedlings. This difference was missing in the *ali1* mutant. Additional expression profiles of *GST6* and *Hsp70* revealed similar results (Supplemental Fig. S3). To comprehensively investigate whether the *ali1* mutant responds to oxo-C14-HSL, we carried out an additional whole transcriptome analysis. We examined differences in the gene expression of plants challenged with 100 nM flg22 after priming with 6 μM oxo-C14-HSL using an RNA sequencing approach. Differentially expressed genes were set as showing a log_2_ fold change of 1 or more and a moderate *p* < 0.05 (Supplemental Datasets S1-S4). Comparison between control and oxo-C14-HSL-pretreated Col-0 seedlings challenged with 100 nM flg22 revealed upregulation of 447 (Supplemental Dataset S1) and downregulation of 29 genes (Supplemental Dataset S2). A similar comparison between control and oxo-C14-HSL-pretreated *ali1* revealed upregulation of 4 (Supplemental Dataset S3) and downregulation of 70 genes (Supplemental Dataset S4) (Fig. 2). Gene Ontology (GO) analysis of Col-0 genes differentially expressed between oxo-C14-HSL-pretreated and control plants, 2 h after the challenge with flg22 showed enrichment of several terms. These are associated with mRNA transcription, movement of cellular components, systemic acquired resistance, cell wall organization and auxin-activated signaling pathway and cellular response (Fig. 2B). The differentially downregulated genes showed enrichment for GO terms associated with abiotic stimulus related to decreased oxygen levels, temperature, salt, osmotic and oxidative stress in the case of Col-0 (Fig. 2B) and glucosinolate metabolism in the case of *ali1*. Taken together, the upregulation of defense-related genes, a typical phenomenon in AHL-primed plants, seems to depend on the presence of ATGALK2/ALI1.

**Figure 2.**
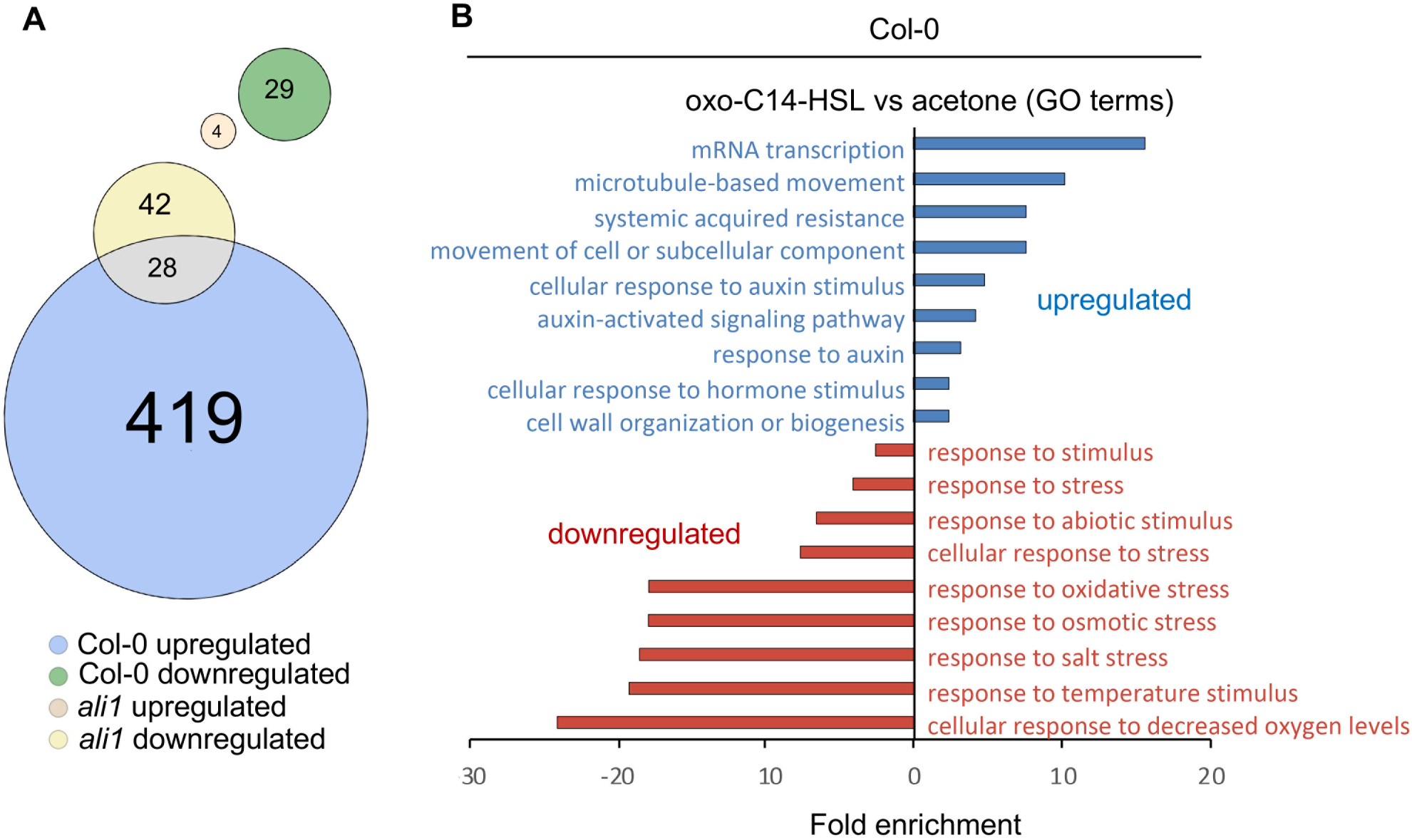
AHL-priming causes differential expression of only few genes in *ali1* mutant. A, Euler diagram of differentially expressed genes after pretreatment with 6 μM oxo-C14-HSL and challenge with 100 nM flg22 for 2 h. The upregulated and downregulated genes from Col-0 are shown in blue and green, respectively whereas the upregulated and downregulated genes from *ali1* are shown in light brown and yellow, respectively. B, The bar plot indicates significantly enriched GO-terms of *Arabidopsis* genes differentially expressed upon oxo-C14-HSL pretreatment and following flg22 challenge, upregulated are shown in blue and downregulated genes in red. Data information: In (A) differentially expressed genes were calculated based on q-value <0.05 and fold change>2 from three independent replica.

To assess whether the absence of the *ATGALK2*/*ALI1* gene has an impact on the oxo-C14-HSL-priming for enhanced resistance, we performed two different *Pseudomonas syringae* pathovar *tomato* (*Pst*)-based pathogen assays. The first assay was performed using the AHL molecule under sterile conditions. *Pst* proliferated on *Arabidopsis* leaves from ca. 10^4^ colony forming units (CFU)/mg FW leaf (2 h after inoculation) to about 10^7^ CFU/mg FW leaf (96 h after the inoculation) (Fig. 3A). However, we observed a reduction in the bacterial CFU number, 96 h after *Pst* inoculation in Col-0 seedlings that were grown on oxo-C14-HSL-supplemented media when compared to control plants. In contrast to the wild type, no enhanced resistance was observed in the *ali1* mutant (Fig. 3A). We performed a similar assay using the oxo-C14-HSL-producing bacterium *Ensifer meliloti expR+* (Hernández-Reyes et al., 2014). In a greenhouse experiment, plants were inoculated three times over three weeks with either 10 mM MgCl_2_ (control), the *E. meliloti attM* strain unable to accumulate the oxo-C14-HSL molecule (bacterial control) or the *E. meliloti expR+* strain. Leaves of Col*-*0 and *ali1* were infiltrated with *Pst* suspension of 2×10^7^ CFU/ml and the presence of *Pst* was monitored for 96 h. A significant reduction of the bacterial count was observed only in leaves of Col*-*0 plants, pretreated with the *E. meliloti expR+* strain (Fig. 3B). This indicates that the AHL-priming for enhanced resistance with oxo-C14-HSL is lost in plants when the *ATGALK2/ALI1* gene is nonfunctional.

**Figure 3.**
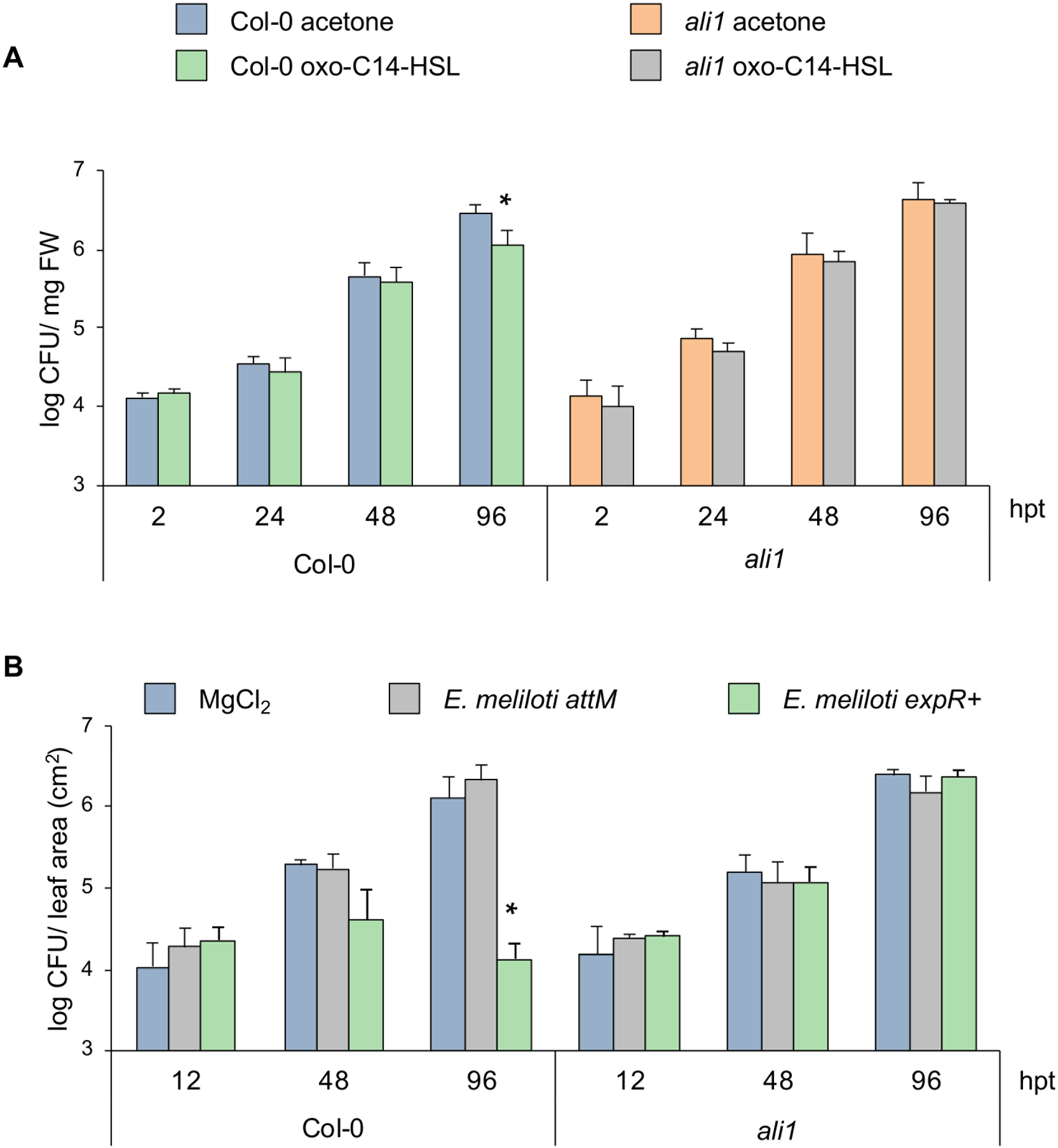
– Oxo-C14-HSL-induced resistance in Arabidopsis against *Pst*, requires the presence of ALI1. A, Pathogenicity assays with *Pseudomonas syringae* pathovar *tomato* (*Pst*) on wild-type Col-0 and *ali1* mutant plants grown for three weeks on 1/2-strength MS containing acetone or 6 μM oxo-C14-HSL and then challenged with *Pst.* The bar represents the mean and error bar represents SD of three replicates. Experiments were performed two times with similar results. B, Pathogenicity assays with *Pst* on wild type Col-0 and *ali1* mutant grown on soil, pre-treated three times with 10 mM MgCl_2_ (control), *Ensifer meliloti attM* or *E*. *meliloti expR*+ and then infiltrated with *Pst*. The bar represents the mean and error bar represents SD of four replicates. Experiments were carried out two times with similar results. Data information: In (A and B) data are presented as mean ± SD. * *P* ≤ 0.05 (Student’s *t*-test).

### YFP-ALI1 is associated with the plasma membrane, tonoplast and endoplasmic reticulum

Next, we examined the cellular localization of ATGALK2/ALI1. To this end, leaves of four-week-old tobacco (*Nicotiana benthamiana*) were infiltrated with a mixture *of Agrobacterium tumefaciens* GV3101 carrying plasmid with sequences of *YFP*-tagged *ATGALK2/ALI1* and LBA4404 carrying plasmid with mCherry-tagged gene sequences of proteins that localizes at diverse subcellular locations (Supplemental Table S2). The results revealed that the localization of the YFP-ALI1 fusion protein resembles the pattern of endoplasmic reticulum (ER) membranes (Fig. 4). Moreover, YFP-ALI1 colocalized with the plasma membrane (PM) and partially with the tonoplast (Fig. 4). Furthermore, YFP-ALI1 did not colocalize with other subcellular organelles including Golgi, nucleolus, plastids and peroxisome (Supplemental Fig. S4) and cell wall as indicated after the plasmolysis (Supplemental Fig. S5). Interestingly, we observed no translocation of the YFP-ALI1 fusion protein after an oxo-C14-HSL treatment (Fig. 4).

**Figure 4.**
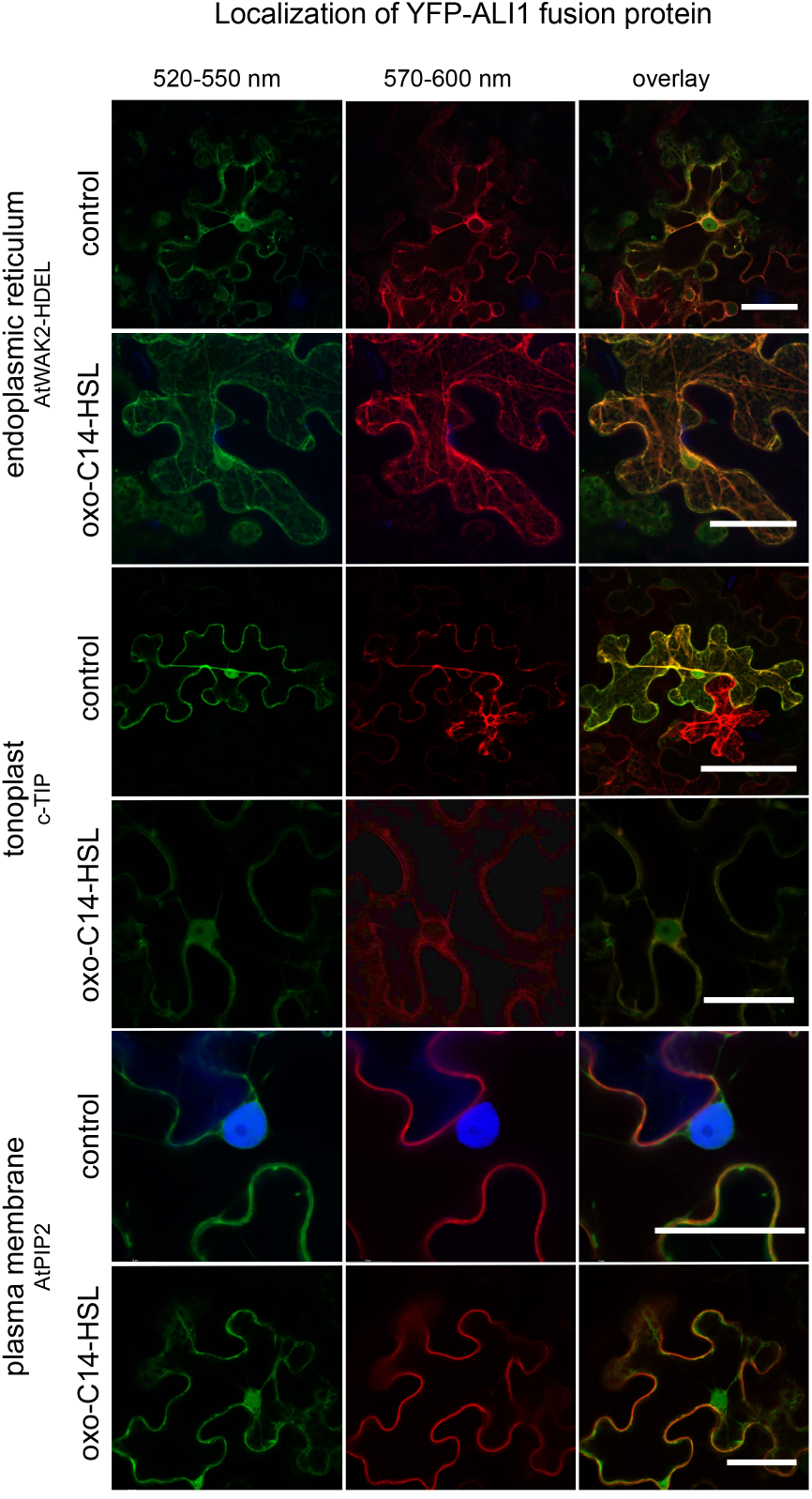
ALI1 protein is localized at the cell’s periphery in PM, ER and tonoplast and do not translocate after oxo-C14-HSL treatment. Confocal images displaying the subcellular localization of YFP-tagged ALI1 before and after oxo-C14-HSL treatment when transiently expressed in cells of *N. benthamiana* leaves. Plasmids carrying YFP-tagged ALI1 version and mCherry-marked proteins localizing in ER (AtWAK2-HEDL), tonoplast (c-TIP) or plasma membrane (AtPIP2) were co-transformed to *N. benthamiana* leaf epidermal cells by *Agrobacterium*-infiltration, and further re-infiltrated after 24 h at the same co-transformed spots of leaf with 6 μM oxo-C14-HSL. The left panel shows in green, fluorescence of YFP-tagged ALI1, whereas the middle panel shows in red, fluorescence of differently localized mCherry-marked proteins (Supplemental Table S2). The right panel shows corresponding merged images, yellow color indicates colocalization. Scale bar = 40 μm.

### Does ALI1 interacts with the bacterial quorum sensing molecule, oxo-C14-HSL?

In order to assess if bacterial quorum sensing molecules interact with plant protein(s), we took advantage of the biotinylated derivative of the *N*-3-oxotetradecanoyl-*L*-homoserine lactone (oxo-C14-HSL) molecule, M4 (Thomanek et al., 2013). We sought to verify the interaction between oxo-C14-HSL and ATGALK2/ALI1 using the fusion 6xHis-ALI1 protein and M4. 6xHis-ALI1 protein was retained on beads bound to M4 but not on beads exposed to the biotin control, suggesting positive interaction between the ALI1 and M4 (Fig. 5A). A 6xHis-tagged Salmonella Plasmid Virulence C (SpvC) protein, a virulence protein in *Salmonella enterica* was used as a negative control in order to verify that M4 does not bind to the 6xHis-tag. Modeling the predicted interaction between ATGALK2/ALI1 and oxo-C14-HSL revealed that the ligand could interact with the central groove of the protein (Fig. 5 B–C and Supplemental Fig. S6).

**Figure 5.**
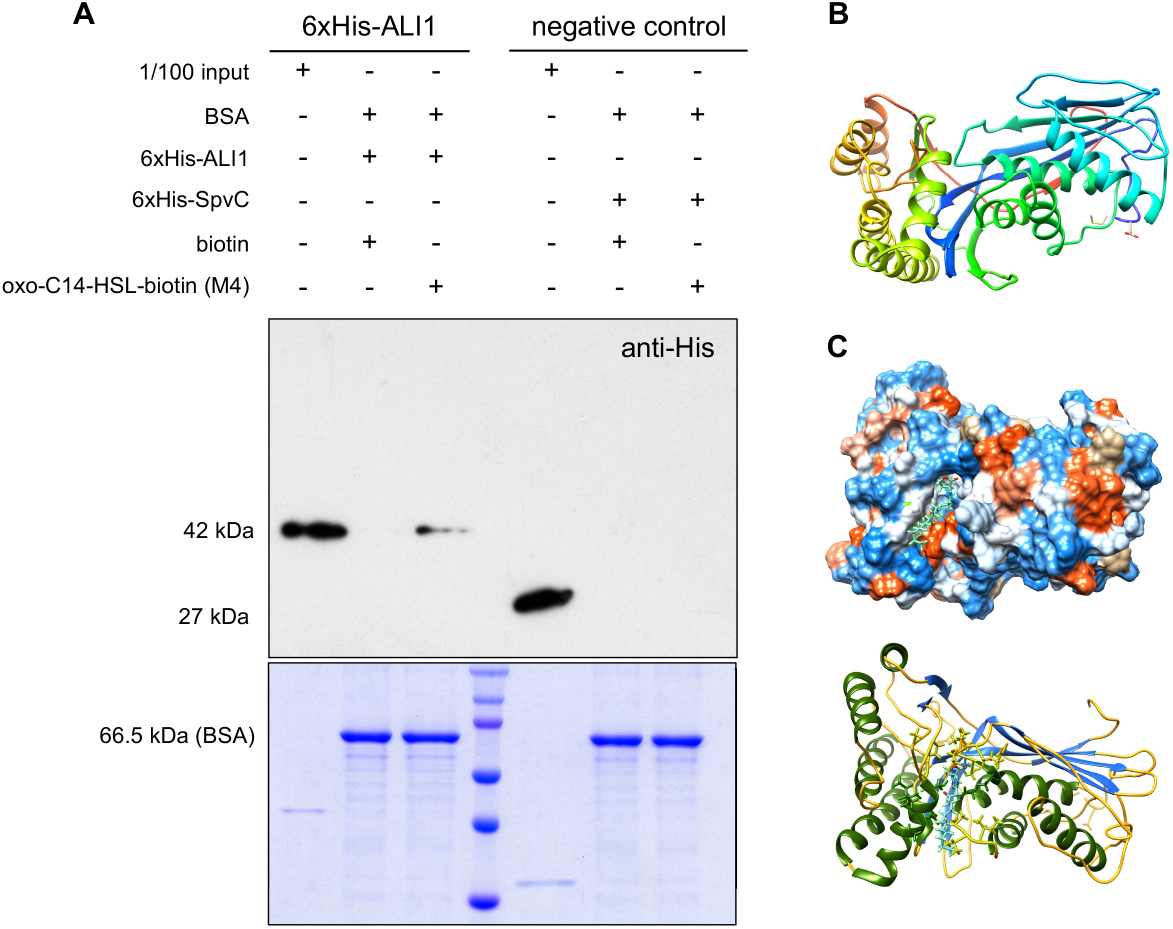
ALI1 interacts with oxo-C14-HSL. A, Pull-down analysis between the His-tagged ALI1 and oxo-C14-HSL (M4) and detection of retained ALI1 on beads bound to M4 but not on beads bound to biotin. Upper panel presents western blot, in which either 1/100 of the input proteins (6xHis-ALI1 or 6xHis-SpvC, as negative control) or proteins resulting from a pull-down were separated, blotted and probed with anti-His antibody. The pull-down setup is indicated above the panel. Pull-down analysis was performed in three independent experiments. Representative blot from those independent experiments is shown. The lower panel presents loading control for the pull-down setup, input shows 1/100 of the protein amount used for the pull-down. B, The tertiary structure of ALI1 protein. (Predicted structure of ALI1 with threading or fold recognition model after docking simulations with oxo-C14-HSL). C, Protein threading or fold recognition predicted model of ALI1 with docked oxo-C14-HSL ligand, simulated in the online docking web server SwissDock. D, Luminescence activity assay indicating the binding of oxo-C14-HSL to ALI1. The binding capacity was assessed by determining the concentration of free oxo-C14-HSL using the *E. coil LuxCDABE* reporter strain after an overnight incubation with 6 nmol ALI1 or LuxR (positive control) and BSA (negative control) with different amounts of oxo-C14-HSL, as indicated.

To further assess the potential interaction between ATGALK2/ALI1 and oxo-C14-HSL, we performed microscale thermophoresis (MST) assay. The combined dose-response curve from both experimental replicates indicated no interaction between ALI1 and oxo-C14-HSL in the analyzed ligand concentration range and under the applied experimental conditions (Supplemental Fig. S7A). One of the reasons for the low chances of observing a binding event could be the distant location of the dye in respect to the protein-ligand binding site. Therefore, we also examined the influence of oxo-C14-HSL on the thermal stability of ATGALK2/ALI1 protein by characterizing its thermal stability and unfolding profile utilizing a nano differential scanning fluorimetry (nanoDSF) assay. For 6xHis-ALI1, two clear unfolding transitions at 50.88°C (inflection point #1) and 66.51°C (inflection point # 2) were observed while for 6xHis-SpvC, one clear unfolding transition at 43.99°C (inflection point #1) was observed (Supplemental Fig. S7B). The presence of oxo-C14-HSL led to a shift of - 1.91 °C (Δ IP #1 column) and −0.88 °C (Δ IP #2 column) in 6xHis-ALI1 and −0.80 °C (Δ IP #1 column) in 6xHis-SpvC (Fig. S7B). Since both proteins (6xHis-ALI1 and 6xHis-SpvC) showed comparable thermal shifts, we concluded that this was probably an artifact. However, thermal shift assays can be biased by unspecific effects such as change in ionic strength, impurities that increase with increasing ligand concentration or other unspecific binding events. Both proteins were shown to be destabilized by a ligand molecule implying that interaction between ALI1 and oxo-C14-HSL was not observed under the applied experimental conditions.

This led us to an indirect method. The concentration of free oxo-C14-HSL was monitored in the presence of both, 6 nmol 6xHis-ALI1 or 6 nmol 6xHis-LuxR (used as a positive control) as well as BSA (used as a negative control). Proteins were incubated overnight with different concentrations of oxo-C14-HSL, and the presence of free molecules was monitored using the reporter *E. coli* strain (*E. coli LuxCDABE*). As revealed, mock and BSA variants induced about 80 to 100% of the possible lux activity (light production capacity) in the reporter *E. coli* strain in the presence of 3 nmol oxo-C14-HSL, indicating that the molecule is freely available. Moreover, 3 nmol oxo-C14-HSL seemed to be sufficient to fully induce the *LuxCDABE* operon. In comparison, the presence of 6 nmol 6xHis-LuxR from *E. meliloti* or 6 nmol of 6xHis-ALI1 from *Arabidopsis* in the system inhibited the induction of the *LuxCDABE* operon significantly. This inhibition was observable until the 9 nmol oxo-C14-HSL variant. In the presence of 6 nmol oxo-C14-HSL (equimolar amount), 50% of the possible *LuxCDABE*-originated luminescence was observed. We concluded, therefore, that LuxR (the native oxo-C14-HSL receptor from *E. meliloti*) and ATGALK2/ALI1 are equivalently able to sequestrate oxo-C14-HSL molecule from the solution (Supplemental Fig. S7C).

### AHL-induced priming is restored in complemented ali1 and ALI1-dependent phenotypes are oxo-C14-HSL specific

In order to assess if the reintroduction of the *ATGALK2/ALI1* gene restores oxo-C14-HSL-dependent phenotypes, we performed gene expression analysis and *Pst* assay using homozygous *ali1* lines expressing Myc-tagged ALI1 under the *35S* promoter (*35S::10xMyc-ALI1*); line #10-2 and #10-19 together with an outcrossed line #10-3 (used as a negative control) (Supplemental Fig. S8). The expression patterns in both complemented #10-2 and #10-19 and the outcrossed (non-complemented) line #10-3, revealed that all seedlings responded to the flg22 challenge (Fig. 6A - C). Furthermore, expression levels of defense-related *WRKY22*, *WRKY29* and *GST6* genes 2 h after flg22 challenge in the complemented #10-19 line, and *WRKY22*, *WRKY29* in the #10-2 line showed higher transcript abundance when pretreated with oxo-C14-HSL (Fig. 6 A – C). This difference was missing in the outcrossed line #10-3. Similarly, the *Pst* assay validated that reintroduction of *ATGALK2/ALI1* gene into the *ali1* mutant, reinstates the ability of AHL-priming. We observed a reduction in the proliferation of *Pst* in oxo-C14-HSL-treated plants compared to control plants in Col-0, 96h after inoculation (Fig. 6D). Furthermore, we observed a similar reduction in *Pst* proliferation in complemented lines, #10-2 and #10-19 after an oxo-C14-HSL pretreatment, indicating enhanced resistance due to AHL-priming (Fig. 6D). This reduction was missing in the non-complemented line #10-3 since no difference in the CFU count between the treatments was observed (Fig. 6D). These results suggest that the reintroduction of *ATGALK2/ALI1* into the *ali1* background restores the ability of AHL-priming.

**Figure 6.**
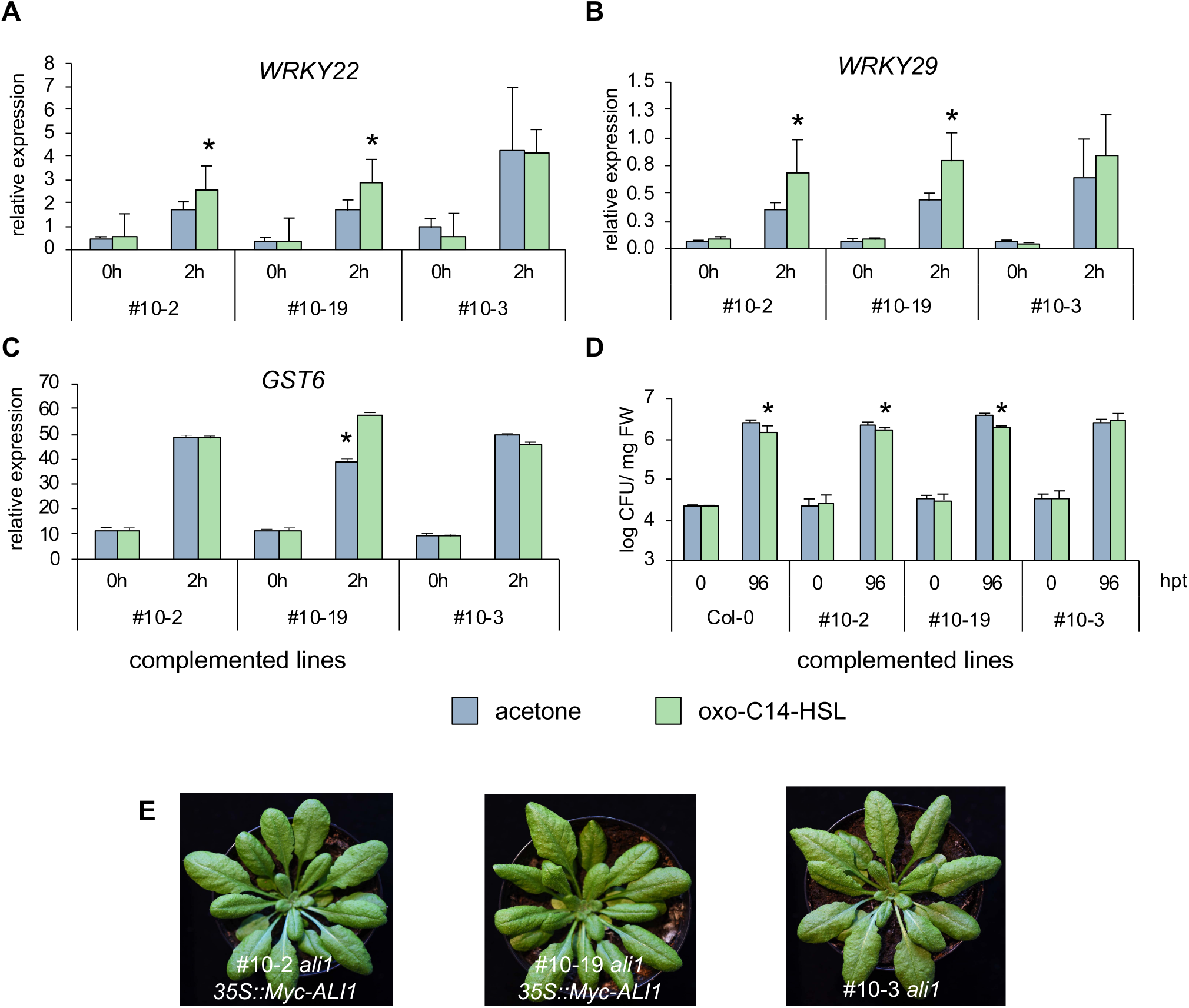
AHL-induced priming is restored in complemented *ali1*. The expression profile of two defense-related genes (A) *WRKY22*, (B) *WRKY29* and (C) *GST6* monitored before (0 h) and after (2 h) 100 nM flg22 challenge in two complemented lines of *ali1:* #10-2 and #10-19 as well as an outcross (non-complemented) line #10-3. The bar represents the mean and error bar represents SD of three replicates. Experiments were carried out two times with similar results. D, Pathogenicity assays with *Pst* on wild-type Col-0, two complemented lines of *ali1:* #10-2 and #10-19 as well as an outcross (non-complemented) line #10-3. Plants grown for three weeks on 1/2-strength MS containing acetone or 6 μM oxo-C14-HSL and then challenged with *Pst.* The bar represents the mean and error bar represents SD of three replicates. Experiments were carried out two times with similar results. E, Representative photographs of six week-old #10-2, #10-19 and #10-3 plants. Data information: In (A - D) data are presented as mean ± SD. * *P* ≤ 0.05 (Student’s *t*-test).

Finally, we aimed to investigate if *ALI1-*dependent phenotypes are specific to oxo-C14-HSL. Our previous reports described AHL-induced growth promotion in plants treated with a short-chained AHL molecule, C6-HSL (Schikora et al., 2011; Shrestha et al., 2020). We performed, therefore, a root growth assay on one-week-old Col-0 and *ali1* seedlings on ½-strength MS agar supplemented with either 6 μM oxo-C14-HSL or 6 μM C6-HSL. Plants were grown vertically to observe the root growth for four weeks. We observed that C6-HSL treatment enhanced root length in Col-0 (Fig. 7A). Moreover, we observed a similar phenomenon of longer roots upon C6-HSL treatment in *ali1* seedlings (Fig. 7A). There was no significant difference in the root length of both Col-0 and *ali1* seedlings between control and oxo-C14-HSL treatments (Fig. 7A). Similarly, we also observed that both Col-0 and *ali1* seedlings growing on plates containing 6 μM C6-HSL had higher biomass (Fig. 7B). Taken together, the influence of short-chained AHL molecules on plant growth does not seem to depend on the presence of functional ATGALK2/ALI1.

**Figure 7.**
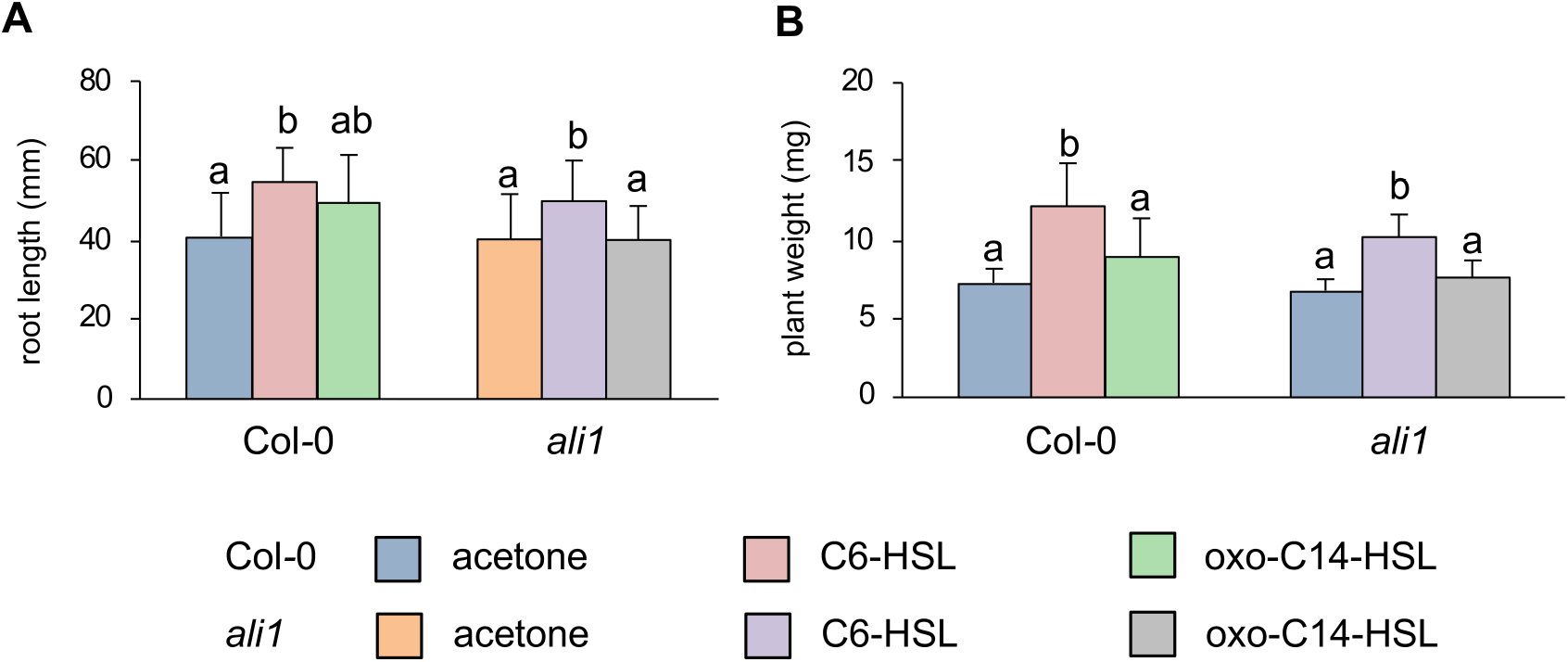
ALI1-dependent phenotypes are oxo-C14-HSL specific. Analysis of (A) root growth and (B) plant weight of Col-0 and *ali1* plants grown for four weeks on 1/2-strength MS containing acetone, 6 μM C6-HSL or 6 μM oxo-C14-HSL (n = 10 plants for each group). Data information: Data are presented as mean ± SD. Different letters indicate *P* ≤ 0.05 (Tukey’s HSD post hoc test). Experiments were carried out three times with similar results.

## Discussion

Quorum sensing (QS) molecules like AHL can modulate interactions between bacteria and eukaryotes. Many studies demonstrated that different AHL molecules have diverse impacts on eukaryotic organisms. Since eukaryotes coevolved with bacteria, it is not surprising that they are able to perceive QS molecules and elicit specific responses. To date, four different studies on mammalian cells have proposed several proteins, which may interact with AHL molecules. Those proteins include Peroxisome Proliferator-Activated Receptors PPARγ and PPARβ (Jahoor et al., 2008), calcoprotein (Seabra et al., 2008), an IQ-motif containing GTPase-activating protein IQGAP1 (Karlsson et al., 2012b) and the bitter receptor T2R38 (Maurer et al., 2015; Gaida et al., 2016). In addition, putative cell-surface or membrane-associated receptors that perceive oxo-C12-HSL were proposed (Shiner et al., 2006; Davis et al., 2010). Until now, there are no studies that have shown a direct interaction of plant protein(s) with QS molecules. Whether the ATGALK2/AHL-Priming Protein 1 (ALI1) interacts with the oxo-C14-HSL molecule could not be sufficiently concluded in this study, we propose however, that ALI1 is required for the oxo-C14-HSL-dependent effects in plants (e.g., AHL-priming).

Thomanek et al. (2013) previously showed that the biotin-tagged oxo-C14-HSL probe was able to interact directly with the bacterial LuxR-type receptor. Here, we used the biotin-tagged probe for pull-down assay. The pull-down assay using a His-tagged ALI1 version suggested that ALI1 indeed interacts with oxo-C14-HSL. Other binding assays like MST assay and nanoDSF assay were however not able to confirm the interaction. This does not eliminate the possible interaction between them since we were able to confirm an oxo-C14-HSL-sequestering ability of ATGALK2/ALI1 through an indirect technique based on the detection of free oxo-C14-HSL by the bacterial biosensor *E. coli LuxCDABE*.

ATGALK2/ALI1 is a putative protein kinase belonging to the GHMP (galactose, homoserine kinase, mevalonate kinase, phosphomevalonate kinase) super kinase family protein (Bork et al., 1993; Holden et al., 2004; Chaves et al., 2009). This protein was predicted to catalyze the phosphorylation of α-D-galactose (Gal) and therefore eliminates the deleterious physiological effect of excess Gal, which often leads to inhibition of root or shoot growth (Yamamoto et al., 1988; Seifert et al., 2002; Warming et al., 2005). A recent report predicted that it might be involved in root and flower development as well as in the abscisic acid signaling pathway and stress response, influencing the expression of various ABA-related genes (Zhao et al., 2013). Nevertheless, based on our experimental evidence, we propose a new function for this protein.

YFP-ALI1 colocalizes with the endoplasmic reticulum (ER), tonoplast and the plasma membrane (PM). Especially, PM and ER contain components that are important for signal transduction and homeostasis regulation. ER-PM contact sites (EPCS) are functional signaling hubs which are formed when ER anchors and couples with PM (Bayer et al., 2017). Since there was no translocation of the ALI1 protein observed after an oxo-C14-HSL treatment, the signal transduction is likely not mediated through translocation but rather through the activity of ALI1 or its interacting proteins. Although we do not know the exact downstream events, several components were already proposed for other AHL molecules.

One of the studies had shown that oxo-C6-HSL and oxo-C8-HSL induced root elongation in *Arabidopsis* requires the G protein-coupled receptors, GCR1 and GPA1 (Liu et al., 2012). Both play a role in root development and are responsive to main plant hormones (Pandey et al., 2006). Furthermore, *Arabidopsis* growth and developmental protein, calmodulin (AtCaM), and the transcription factor AtMYB44, are involved in oxo-C6-HSL-induced root elongation (Zhao et al., 2015; Zhao et al., 2016). However, those specific components are likely to be involved in response to a particular, in this case, short-chained AHL molecule(s) and not required for ALI1-mediated responses to oxo-C14-HSL. Previous reports suggested that the enhanced and prolonged activation of the MAP kinases, MPK3 and MPK6, along with the upregulation of defense-related genes are associated with oxo-C14-HSL-mediated priming for enhanced defense response in plants (Beckers et al., 2009; Schikora et al., 2011; Schenk and Schikora, 2014; Shrestha et al., 2019). Our current observations are consistent with these studies. The *ali1* mutant showed neither an increase in kinase activity nor an enhanced expression of defense-related genes, both hallmarks of AHL-priming. Similarly, the pathogenicity assay with *P*. *syringae* indicated that the AHL-priming for enhanced resistance is absent in plants without functional *ATGALK2/ALI1*. However, this phenomenon is restored once the *ALI1* gene is reintroduced into the *ali1* mutant.

Taken together, we present here several levels of confirmation that ATGALK2/ALI1 is required for the oxo-C14-HSL-induced primed state in *Arabidopsis thaliana*. Studies focusing on the identification of AHL-priming mechanisms are necessary, since they open new opportunities for the application of AHL molecules in agriculture.

## Materials and methods

### Plant material and growth

Seeds from wild-type *Arabidopsis thaliana* Col*-*0 and the *ali1* (N560407; *At5g14470*) mutant, obtained from the Salk Institute, as well as complemented (pGWB21; *35S:: 10xMyc-ALI1*) and outcrossed lines were used in the study. Seed sterilization and Arabidopsis growth was carried out as was described in Shrestha et al. (2020). For MAP kinase assay, gene expression analysis and RNA sequencing approaches, two-week old seedlings were transferred to six-well plates with 3 ml 1/2-strength MS medium per well prior to *N*-acyl homoserine lactone (AHL) pretreatment. Root growth assay and sterile *Pseudomonas syringae* pathogenicity assay were performed as described in Shrestha et al. (2020) except for AHL molecules used. For non-sterile *P. syringae* pathogenicity assay and *ali1* mutant complementation, two-week old seedlings were transferred to pots with standard bedding soil (Fruhstorfer Erde: Perlite (1:1)) and grown under short-day conditions at 8/16 h (day/night) photoperiod at 22°C for another three weeks in the greenhouse. In order to induce flowering, six-week old plants were grown under long-day conditions (day/night 16/8 h and 22°C photoperiod, light intensity of 150 μmol/m^2^s and 60% humidity).

### ALI1 expression and purification

Gateway destination vector pDEST17 subcloned with *ALI1* ORF was transformed into *E. coli* BL21 competent cells (New England Biolabs). For detailed information, see Supplemental Methods.

### Pull-down and binding assays

The interaction of ALI1 protein with oxo-C14-HSL was verified by pull-down assay. 6xHis-tagged ALI1 and 6xHis-tagged control protein were incubated with streptavidin beads (Sepharose^®^ Bead Conjugate, Cell Signaling Technology) that were already loaded with M4 (oxo-C14-HSL tagged with biotin) or biotin control and incubated overnight before washing with washing buffer. The final bead-bound protein fractions were eluted with Laemmli buffer (62.5 mM Tris-HCl (pH 6.8), 2% SDS, 10% glycerol, 0.01% bromophenol blue) containing 2% β-mercaptoethanol and subjected to western blotting.

30 μl of streptavidin beads were blocked using blocking buffer (0.05% (w/v) BSA in 100 mM phosphate buffer at pH 7.4) for half an hour at RT and further loaded with 60 mM biotin or 60 mM biotin-labeled oxo-C14-HSL (M4) for 30 mins and centrifuged at 67 *g* for one min. After removal of supernatant, 100 μg 6xHis-tagged proteins were added and incubated overnight at constant stirring at 4°C. After the beads were washed five times with blocking buffer, bound proteins were eluted by adding 40 μl of Laemmli buffer and heated at 95°C for 5 min. 20 μl of eluted protein samples were separated on SDS gel (12%) and transferred to PVDF membrane (Immun-Blot^®^ PVDF, Bio-Rad) through semi-wet blotting protocol. The membranes were blocked with 5% fat-free milk and thereafter probed with primary antibodies 6xHis Tag Monoclonal Antibody (Invitrogen) followed by incubation with horseradish peroxidase-labeled secondary antibody. Blots were developed using ServaLight chemiluminescent substrate (SERVA).

In order to verify the binding capacity of ALI1 to oxo-C14-HSL, we used an additional indirect technique based on the detection of free oxo-C14-HSL by the bacterial biosensor *E. coil LuxCDABE*. We used BSA as a negative control and the *E. meliloti*-originated LuxR receptor as purified His- and GST-versions, as positive control, and the ALI1, purified as GST-ALI1 and 6xHis-ALI1 fusion proteins. Six nmol of each protein was used. The oxo-C14-HSL was added in a range from 0 to 18 nmol and incubated with proteins overnight. Unbound oxo-C14-HSL was detected after overnight incubation using the bacterial biosensor *E. coli LuxCDABE,* as described in Schenk et al. (2014).

### Pretreatment with AHL

Pretreatment with AHL was conducted as in Shrestha *et al*, (2020) except for the AHL molecules used. *N*-hexanoyl-*L*-homoserine lactone (C6-HSL) and *N*-3-oxotetradecanoyl-*L*-homoserine lactone (oxo-C14-HSL) (Sigma-Aldrich) at concentrations as indicated were used in this study.

### MAP kinase activity assay

Arabidopsis seedlings Col-0 and *ali1* (N560407) mutant were pretreated with 6 μM oxo-C14-HSL or acetone for 3 days and collected 0, 30, 60 or 120 min after eliciting with 100 nM flg22 peptide. Protein extraction from Arabidopsis seedlings and western blotting were performed as in Shrestha et al. (2019).

### Gene expression analysis

Wild-type Col-0 and *ali1* (N560407) Arabidopsis seedlings pretreated with oxo-C14-HSL were harvested at 0, 2, 4, 6, 24 and 48 h post 100 nM flg22 treatment. Extraction of RNA and synthesis of cDNA was carried out as described in Shrestha et al. (2020). Quantitative RT-PCR (qPCR) was performed using primers listed in Supplemental Table S1. All expression levels were normalized to the expression of *Ubiquitin ligase* (*At5g25760*). The experiments were performed in four independent replications.

### Challenge with *Pseudomonas syringae*

*Pseudomonas syringae* pv. *tomato* DC3000 (*Pst*) pathogen assay on sterile Arabidopsis was performed as in Shrestha et al. (2020). The bacterial culture was adjusted to OD_600_ = 0.1. For *Pst* challenge on non-sterile condition, three days after the last inoculation with *E. meliloti* or MgCl_2_, four leaves from each treated plant were infiltrated with *Pst* inoculation solution adjusted to OD_600_ = 0.02. The leaves were infiltrated with *Pst* using a needleless syringe and four-leaf discs were collected from two leaves at 12, 48 and 96 h post infiltration. The leaf discs were homogenized in 10 mM MgCl_2_ and subsequently diluted. Duplicates of the dilution were plated on King’s B agar plates containing selective antibiotics to assess the colony-forming units (CFU) number. The experiments were performed in three independent replications.

### Assessment of localization of ALI1

In order to assess the subcellular localization of ALI1, *ALI1* ORF was subcloned into pGWB441 vector containing yellow fluorescent protein (YFP) gene and the Cauliflower Mosaic Virus (CaMV) 35S promoter and transformed into NEB 5-alpha Competent *E. coli* (New England Biolabs). Cloned vector extracted from transformed *E. coli* DH5α was used to transform competent *Agrobacterium tumefaciens* GV3101. YFP-tagged *ALI1* in *A. tumefaciens* GV3101 along with several *A. tumefaciens* LBA4404 containing mCherry-tagged gene sequence of proteins (Supplemental Table S2) that localizes at different subcellular organelles were grown overnight in 5 ml LB liquid medium with selective antibiotics at 28°C. The bacterial cultures were centrifuged at 2500 *g* for 10 min and resuspended with infiltration buffer at final OD_600_ of 0.05 and infiltrated into leaves of four-week old tobacco (*N. benthamiana*) plants. Equal OD_600_ 0.05 of *A. tumefaciens* GV3101 containing YFP-tagged *ALI1* and *A. tumefaciens* LBA4404 containing mCherry-tagged gene sequence of a protein that localizes at a subcellular organelle were mixed in infiltration buffer and infiltrated on the abaxial side of the leaves with 1 ml needleless syringe. The infiltrated leaves were visualized for protein localization on confocal laser scanning microscopy (CLSM) after two to three days after infiltration. Samples were analyzed using confocal laser scanning microscopy, YFP was observed using 514 nm Ex and 520 – 550 nm Em range, and mCherry using 560 Ex and 570-600 Em settings.

### Complementation of *ali1* mutant

For complementation of the *ali1* mutant (N560407), pGWB21 (35S, N-ter 10xMyc) destination vector was used. For detailed information, see Supplemental Methods.

### Statistical analysis

All experiments were performed with at least three independent biological replicates. The GENMOD procedure from SAS 9.4 (SAS Institute Inc., Cary, NC, United States) was used for the analysis of variance. For multiple comparisons in root growth assay, the *p*-value was adjusted by the method of Tukey’s honestly significant difference (HSD) post hoc test. The class variable was treatment (C6-HSL, oxo-C14-HSL and acetone control). Quantitative PCR assays were performed in four biologically independent experiments and *Pst* assays were performed in three biologically independent experiments. *p* values < 0.05 in the Student’s *t*-test were considered as indicative for a significant difference. Western blot analysis was performed in three independent experiments, representative blot is shown.

## Acknowledgements

The authors thank Wolfgang Maison, Annette Niehl, Hendrik Reuper, Khalid Amari Baba, Davide Faggionato, Svenja Lindenau and Miriam Breuer for their help in providing us with materials to perform some of the experiments. We would also like to thank Dr. Thomas Schubert and 2bind GmbH for their generous support in performing MST binding assay and nanoDSF assay. The work of Abhishek Shrestha was supported by DFG, grant nr. SCHI 1132-11 granted to Adam Schikora. The work of Casandra Hernandez-Reyes was supported by CONACYT.

## Author contributions

AbS, CSR, STS and AS conceived the research project. AbS performed ALI1 purification, pull-down assay, assessment of MAP kinase activity, western blot analysis, CLSM images, complementation of *ali1* mutant, in silico screening for docking, transcriptome analysis and analyzed the data. AbS, MG, JK and CSR performed cloning and AbS and MG performed RNA extraction and gene expression analysis and *Pst* pathogen assay. AbS and AS wrote the manuscript and all authors contributed writing, reviewing and editing the manuscript.

