## Supplementary Material for "Novel piece of the puzzle: ALI1 is required for oxo-C14-HSL priming in Arabidopsis"

### Supplemental Methods

**In Silico screening for docking of oxo-C14-HSL against ALI1 (*At5g14470*).** The sequence of the ALI1 (ATGALK2) protein was obtained from TAIR and the best predicted model of ALI1 was generated using i-TASSER server https://zhanglab.ccmb.med.umich.edu/I-TASSER/ (Yang and Zhang, 2015). Similarly, 3D conformer of the ligand oxo-C14-HSL was retrieved from PubChem Structure (Kim et al., 2016). The 3D structure of both protein and ligand was visualized using UCSF Chimera (Pettersen et al., 2004) where the model was prepared for docking in order to attain a clean protein model for further analysis. Docking simulations of ALI1 protein and oxo-C14-HSL ligand was performed in SwissDock (http://www.swissdock.ch/docking) (Grosdidier et al., 2011), a web server that provides prediction of molecular interaction between the ligand and the target protein. The ‘docking type’ was set to accurate type and ‘Definition of the region of interest’ was set to default and so was ‘flexibility’ which allows flexibility for side chains within 0 Å of any atom of the ligand in its reference binding mode. The binding modes were scored using their FullFitness and clustered, of which they were ranked on the basis of the average FullFitness of their elements (Grosdidier et al., 2007). The prediction file from SwissDock was further inspected in UCSF Chimera and the chimera model with the lowest energy was selected for the identification of amino acid residues that were at a distance of less than 5 Å from each atom of the ligand.

**Cloning approach.** *ALI1* ORF sequence (*At5g14470*) was amplified from cDNA of Col-0 Arabidopsis using ALI1 specific primers (Supplemental Table S1). Subsequently, *ALI1* ORF sequence was supplemented with *attB* sites using sequence-specific primers containing *attB* sites (Supplemental Table S1). Thereafter, the PCR product was purified using PEG precipitation protocol (Schmitz and Riesner, 2006). BP clonase recombination reaction was performed to subclone *ALI1* into the pDONR207 vector using Gateway® BP Clonase™ II Enzyme Mix following the manufacturer’s protocol. NEB 5-alpha Competent *E.* *coli* (New England Biolabs) cells were transformed with the product of recombination reaction using the heat shock method. Colony PCR was performed using DNR3 and DNR5 primers and amplicons were digested with BamHI and ran on a gel for further analysis. Positive constructs were sent for sequencing. LR clonase recombination reaction was performed to insert *ALI1* ORF into different destination vectors (pGWB21 or pGWB441) using Gateway™ LR Clonase™ II Enzyme Mix following the manufacturer’s protocol. Different destination vectors were used for the study. For protein expression and purification, pDEST17 was used whereas pGWB441 was used for cellular localization studies.

**Assessment of *ali1* mutant.** Leaves were collected from three-week old wild type Col*-*0, *ali1* and a random T-DNA insertion mutant that was used as T-DNA insertion mutant control. DNA extraction of the leaf material was performed using DNeasy® Plant Mini Kit (Qiagen) following the manufacturer’s protocol. DNA concentration and quality were measured using the Nanodrop Bioanalyzer. PCR reactions were carried out for both T-DNA insertion and for gene sequence using two sets of primer pairs (Supplemental Table S1) and the PCR amplicons were subjected to agarose gel.

**cDNA library construction, sequence processing and transcriptome analysis.** We performed transcriptome analysis of Col-0 and *ali1* seedlings that were pretreated for three days with 6 µM oxo-C14-HSL or acetone control and subsequently elicited with 100 nM flg22 in order to determine the differences in their defense responses. All treatments were performed in triplicates. RNA was isolated from Arabidopsis seedlings using RNeasy Plant Mini Kit (Qiagen) according to the manufacturer’s recommendations. One μg of total RNA was taken for DNAse digestion using the PerfeCTa DNAse I (Quanta Biosciences) and subsequently cDNA synthesis was carried out using the qScript cDNA Synthesis kit (Quanta Biosciences) according to the manufacturer’s recommendations. Library construction and sequencing were performed on BGISEQ-500 (BGI Tech Solutions, Hong Kong). Raw sequencing reads were cleaned by removing adaptor sequences, reads containing poly N-sequences and low-quality reads. The cleaned sequence reads (100 bp single end) were analyzed using RNAStar (Version 2.4.0d-2) (Dobin et al., 2013), cufflinks (Version 2.2.1.0), cuffmerge (Version 2.2.1.0) and cuffdiff (Version 2.2.1.5) (Trapnell et al., 2010). The FPKM (Fragments Per Kilobase Million) and the significant differences between transcriptional profiles of Arabidopsis seedlings Col-0 and *ali1* between the treatments were calculated based on q-value <0.05 and fold change >2 (Supplemental Dataset S1). Significantly enriched genes related to specific GOs (Gene Ontology) were identified using web-based tool PANTHER (http://geneontology.org) using the PANTHER Overrepresentation Test (Released 2020-02-14), *Arabidopsis thaliana* (TAIR), GO biological process complete and Fisher’s exact test with false discovery rate (FDR). The raw data were uploaded to GEO NCBI Sequence Read Archive with the number GSE156726. The Euler diagram was created using the R package ‘eulerr’ (Larsson, 2020).

**Characterization of *ali1* mutant.** For genotyping of the T-DNA insertion *ali1* mutants*,* PCR based screening was performed for knockout mutation*.* Genomic DNA was extracted from wild type Col-0, mutant *ali1* and T-DNA insertion mutant that were growing on soil for 3 weeks using Qiagen DNeasy Plant Mini Kit following the manufacturer’s protocol. The first PCR was performed with left border primer LBb1-3 and gene specific forward primer (Supplemental Table S1). The band was observed only in *ali1* mutant and not in the other T-DNA insertion mutant and wild-type Col-0 indicating that T-DNA was indeed inserted in *ALI1* locus (Supplemental Fig. S3). Similarly, the second PCR that was performed with *ALI1* gene specific primers (Supplemental Table S1) resulted in the band in wild-type Col-0 and T-DNA insertion mutant but not in *ali1* mutant validating that there is a modification in the gene (Supplemental Fig. S3). Furthermore, third PCR was carried out with *ALI1* cDNA specific primers (Supplemental Table S1) to verify whether the transcription of *ALI1* gene was impeded or not. The band was seen in cDNA of Col-0 pretreated with both oxo-C14-HSL and acetone control but not for *ali1*.

**Inoculation with *Ensifer meliloti* strains.** *Ensifer meliloti* Rm2011 *expR*+ (M. McIntosh) and *E. meliloti* Rm2011 (pBBR2-attM) carrying the lactonase gene *attM* from *Agrobacterium tumefaciens* (Zarkani et al., 2013) were grown in Tryptone Yeast extract (TY) medium until the OD_600_ reached 0.6 - 0.8. Bacterial cultures were centrifuged at 2500 *g* for 10 min and resuspended in 10 mM MgCl_2_. The rhizosphere of Arabidopsis was inoculated three times over three weeks with 10 ml of OD_600_ = 0.1 using: *E. meliloti expR*+, *E. meliloti* *attM* culture solution or 10 mM MgCl_2_ as control. The production of AHL on the root surface as well as establishment of the bacteria was previously demonstrated by (Zarkani et al., 2013).

**Localization of ALI1 upon oxo-C14-HSL treatment.** In order to assess a possible relocalization of ALI1 upon oxo-C14-HSL treatment, the leaves that were pre-infiltrated for 24 h with *A. tumefaciens* mix solution were infiltrated again with 6 µM oxo-C14-HSL. Similarly, acetone on infiltration buffer was used as a control. The infiltrated leaves were visualized for ALI1 localization 1 and 2 days after the infiltration with oxo-C14-HSL.

**Complementation of *ali1* mutant.** For complementation of the *ali1* mutant (N560407), pGWB21 (35S, N-ter 10xMyc) destination vector was used. Competent *E. coli* DH5α was transformed with the product of LR recombination reaction using heat shock method. Colony PCR was performed using vector and gene specific primers (Supplemental Table S1) and obtained constructs were verified via sequencing. *A. tumefaciens* strain GV3101 was transformed with the desired vector constructs by electroporation and selection of the positive colonies was performed through colony PCR using vector and gene specific primers (Supplemental Table S1). The positive cells were used to transform the *ali1* mutant using the floral-dip method. Seeds of the *ali1* mutant were first surface sterilized and grown on ½-strength MS media plates for two weeks and later transferred on standard bedding soil (Fruhstorfer erde: Perlite (1:1)). The two-week old seedlings were grown on controlled condition (day/night 8/16 h and 22°C photoperiod, light intensity of 150 µmol/m^2^s and 60% humidity) in a growth chamber for two months before transferring them to the greenhouse condition (day/night 16/8 h and 22°C photoperiod, light intensity of 150 µmol/m^2^s and 60% humidity) until flowering. *A. tumefaciens* GV3101 strains containing *ALI1* ORF in pGWB21 was grown overnight in LB medium containing selective antibiotics. 250 µl of pre-culture was inoculated on 250 ml LB medium containing selective antibiotics and was grown overnight at 28°C until the O.D_600_ reached 1. The culture was collected and centrifuged at 2500 *g* for 10 min at RT. The pellets were resuspended in 250 ml of 5% sucrose solution and Silwet L-77 was added at a concentration of 0.02%. Floral-dip method (Clough and Bent, 1998) was employed to stably transform the *ali1* mutant. Inflorescences were dipped into the *Agrobacterium* suspension for 30 seconds. The plants were then covered with a hood for 48 h and left to grow in the greenhouse under long day conditions. Watering of the plants was stopped after the first pods began to dry. Seeds were harvested after complete drying of the inflorescences and seeds were selected on selective ½-strength MS medium plates. The seedlings growing on selective plates were transferred on soil pots and later genotyped in order to identify the positive heterozygous complemented lines (T_0_). The positive complemented lines were self-pollinated and the generated seeds (F_1_) were checked for homozygosity using gene-specific and tag-specific primers (Supplemental Table S1).

**Western blot on complemented *ali1* lines.** Arabidopsis seedlings Col*-*0, complemented *ali1* (N560407) mutants #10-2 and #10-19 and outcross line #10-3 were first grown on ½-strength MS plates for two weeks and then transferred on soil pots. The plants were grown for additional four weeks and one leaf was collected and homogenized. Proteins were extracted from homogenized plant samples using Laemmli buffer (62.5 mM Tris-HCl (pH 6.8), 2% SDS, 10% glycerol, 0.01% bromophenol blue) further supplemented with Triton-X (10%) and additional SDS (4%). The homogenized plant samples were vortexed vigorously, subsequently cooked at 95°C for 10 mins and briefly centrifuged. Fifteen μl of total protein was run on SDS gel (12%) and transferred to PVDF membrane through semi-wet blotting protocol. The membranes were blocked with 5% w/v fat-free milk and thereafter probed with primary antibody Myc-tag antibody (Chromotek GmbH), followed by incubation with horseradish peroxidase-labeled secondary antibody Anti-rat IgG (Cell Signaling Technology). Blots were developed using chemiluminescent substrate (ServaLight Vega Luminol solution, SERVA).

**ALI1 expression and purification.** Gateway destination vector pDEST17 subcloned with *ALI1* open reading frame (ORF) was transformed into *E.* *coli* BL21 competent cells (New England Biolabs). Colonies were screened and selected with gene-specific and vector-specific primers (Supplemental Table S1). Positive colonies were grown in 50 ml of LB containing 100 mg/ml ampicillin at 37°C until OD_600_ reached 0.6. The expression of the protein was induced by adding 1 mM isopropyl β-D-1-thiogalactopyranoside (IPTG) overnight at room temperature (RT). All further steps were carried out at 4°C or on ice. Bacterial cells were collected and centrifuged at 7379 *g* for 15 min and lysed through sonication with ice-cold lysis buffer (50 mM Tris-HCl, 300 mM NaCl, 0.1 % Triton-X, DNase, lysozyme supplemented with protease inhibitor (Roche) and thereafter centrifuged at 14462 *g* for 10 minutes. The resulting supernatant was added to a solution containing nickel resin beads (HisPur^TM^ Ni-NTA Resin, Thermo Fisher Scientific) and binding buffer (50 mM sodium phosphate buffer at pH 8, 500 mM NaCl, 10 mM imidazole, 2.5% glycerol) for 30 min with continuous rotation. Lysate-bead mix was subsequently transferred to column and washed 5 times with washing buffer (50 mM sodium phosphate buffer at pH 8, 500 mM NaCl, 20 mM imidazole, 2.5% glycerol) and eluted with elution buffer (50 mM sodium phosphate buffer at pH 8, 500 mM NaCl, 250 mM imidazole, 2.5% glycerol). The concentration of 6xHis-tagged proteins was quantified using Bradford assay (Roti-Quant, Roth). *E. coli* BL21 transformed with pDEST17-SpvC was used as a 6xHis-tagged protein control.

**Microscale thermophoresis (MST) binding assay.** Preceding to MST bioassay, both proteins (6xHis-ALI1 and 6xHis-SpvC used as His-tagged protein control) were labelled with Red-Tris-NTA fluorescent dye and diluted into the labeling buffer (50 mM phosphate buffer at pH 8.0, 150 mM NaCl, 0.005% Tween-20 and 0.16% acetone) to reach the labeling concentration of 0.2 µM. After labeling, dye removal was not required as all dye molecules were bound by a protein due to labeling stoichiometry and therefore, no free dye was present in the labeling batch. The final concentration of total protein and fluorescence was 200 nM and 100 nM respectively. Premium-coated capillaries were chosen for further experiments as no adhesion of both labeled proteins was observed. The intrinsic MST noise of both labeled proteins was acceptably low (<5.0 units) under the applied experimental parameters and the used assay buffer. MST binding assay was performed on Monolith NT.115 Pico (red-nano) at 25°C, with 90% LED power and 40% laser power. 5 µl of the fluorescent target proteins were used at constant 50 nM where 5 µl of the ligand oxo-C14-HSL was titrated from 100 µM down in 16 (1:1 dilution) steps. To ensure reproducibility, measurements were taken repeatedly on two independent experiments. The data were analyzed by ligand concentration (mol/l) against normalized fluorescence of labelled proteins. Curve fitting were performed by using the K_D_ fit derived from the law of mass action according to the binding model, the affinity is calculated and stated as EC50 or K_D_ values. The amplitude of the binding curve was assessed and signal to noise ratio which is amplitude divided by noise. Noise is standard deviation of difference between experimental data and fitted data.

**Nano-differential scanning fluorimetry (nanoDSF) assay.** Prior to nanoDSF assay, both protein samples (6xHis-ALI1 and 6xHis-SpvC used as His-tagged protein control) and ligand oxo-C14-HSL were diluted in assay buffer (50 mM phosphate buffer at pH 8.0, 150 mM NaCl, 0.005% Tween-20 and 0.167% acetone) to have a final concentration of 5 µM and 100 µM respectively. The experiments were performed on a Prometheus NT.48 device equipped with additional back reflection options for detection of target protein aggregation via light scattering method. The temperature of the assay ranged from 20°C to 95°C with heating speed of 1°C/min. The analysis method of ratio 350/330 nm for protein unfolding (Tm) was used and data were analyzed using the PR.StabilityAnalysissoftware from Nanotemper Technologies. The thermal denaturation of the target proteins were monitored via its intrinsic tryptophan fluorescence. Thermal shift was calculated by assessing difference of melting temperatures (Tm) between ligand-bound and apo-state of a protein.

### Supplemental Figures

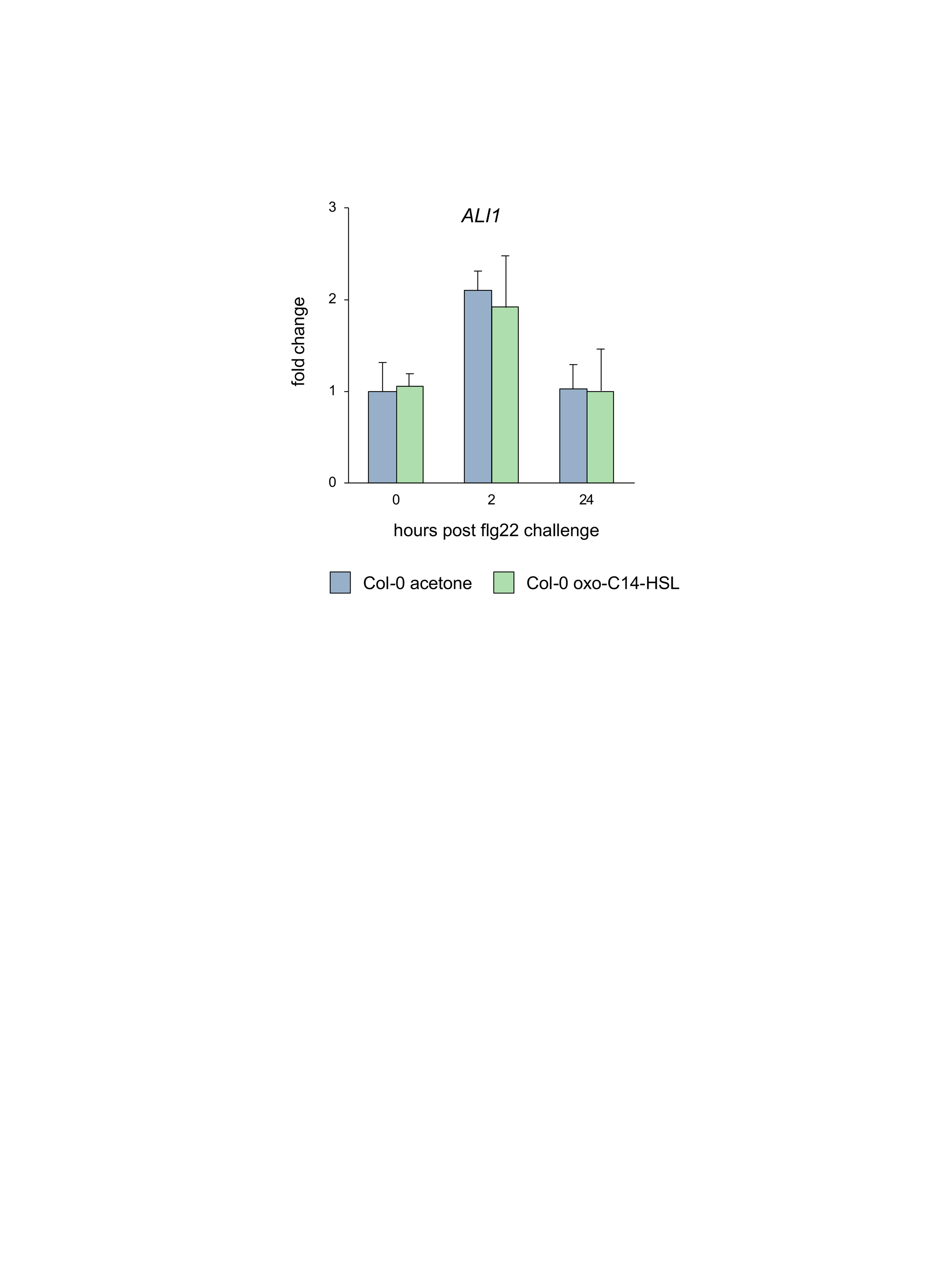

#### Supplemental Figure S1. The expression of *ALI1* is not influenced by oxo-C14-HSL.

Expression profile of *ALI1* was monitored at three time points after challenge with 100 nM flg22 in *Arabidopsis* wild-type Col-0. Plants were grown on sterile hydroponic system and pretreated with 6 µM oxo-C14-HSL or acetone (solvent control) three days prior to flg22 challenge. The abundance of each gene transcript was normalized with *Ubiquitin ligase* (*At5g25760*) transcript and 0 hpt (hours post treatment) levels. The bar represents mean and error bars SD from four independent biological replicates.

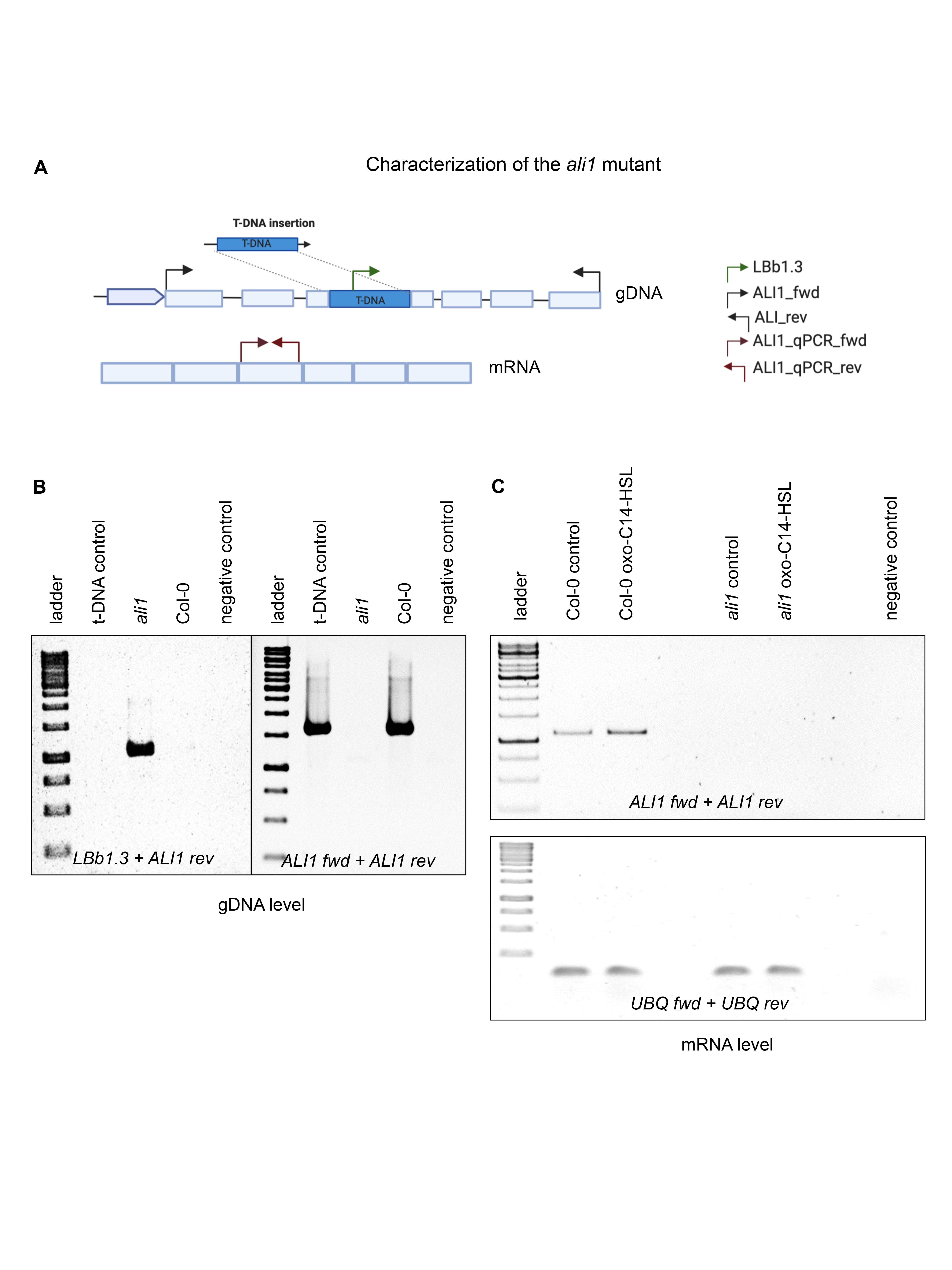

#### Supplemental Figure S2. The *ali1* mutant has a T-DNA insertion in *At5g14470* position and is homozygous.

A, Graphical representation of T-DNA insertion in *ALI1* gene, indicating primers (Supplemental Table S1) used for characterization of the mutant. B, PCR-based characterization of the genetical structure of *ali1*. The insertion of T-DNA and the absence of functional *ALI1* gene in *ali1* mutant however, absence of T-DNA insertion and the presence of functional *ALI1* gene in wild-type Col-0 were evidenced by PCR products using primers as indicated. C, The expression of *ALI1* in both acetone and oxo-C14-HSL-pretreated Col-0 plants as well as the absence of *ALI1* expression in both acetone and oxo-C14-HSL-pretreated *ali1* mutant, as evidenced by PCR approach. The expression of housekeeping gene *Ubiquitin* *ligase* (*UBQ*) in both acetone and oxo-C14-HSL-pretreated Col-0 and *ali1* mutant. T-DNA left border-specific LBb1-3 and *ALI1-*specific reverse primers were used to detect the presence of T-DNA insertion in *ALI1* gene, whereas gene-specific primers, ALI1 fwd and ALI1 rev were used in a separate reaction to detect the presence of *ALI1* gene in both, DNA and mRNA levels and *UBQ* gene-specific primers, UBQ fwd and UBQ rev were used in a separate reaction for the internal control.

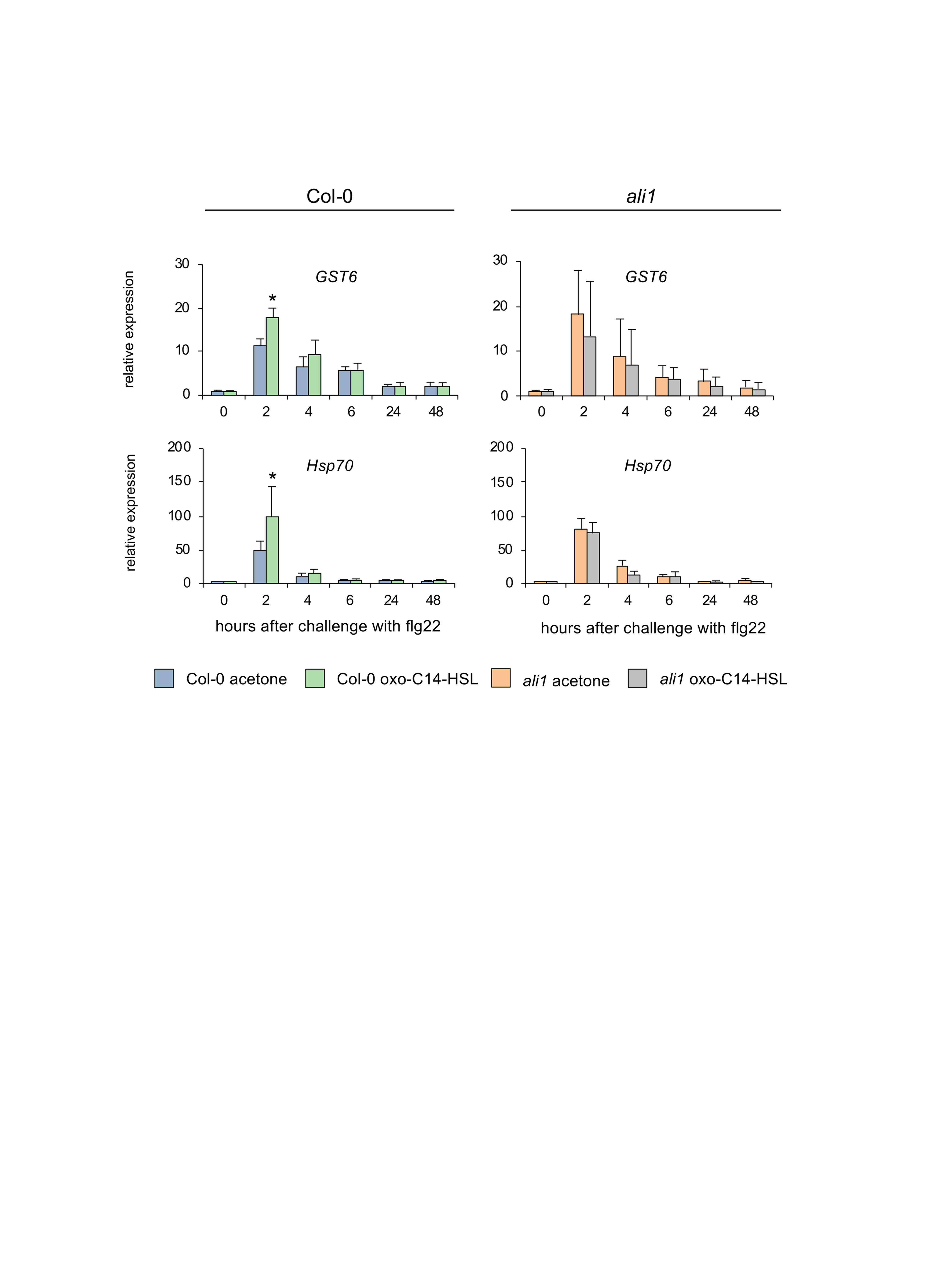

#### Supplemental Figure S3. Enhanced activation of defense-related genes due to AHL-priming is missing in *ali1*.

Expression profile of defense-related genes *GST6* and *Hsp70* was monitored at various time points after 100 nM flg22 challenge in Col-0 and *ali1* mutant. Plants were grown on a sterile hydroponic system and pretreated with 6 µM oxo-C14-HSL or acetone (solvent control) for three days prior to flg22 challenge. The abundance of each gene transcript was normalized with *Ubiquitin* *ligase* (*At5g25760*) transcript and 0 hpt (hours post treatment) levels. Error bars represent SD from four independent biological repetitions. * indicates *p* < 0.05 in Student’s *t*-test.

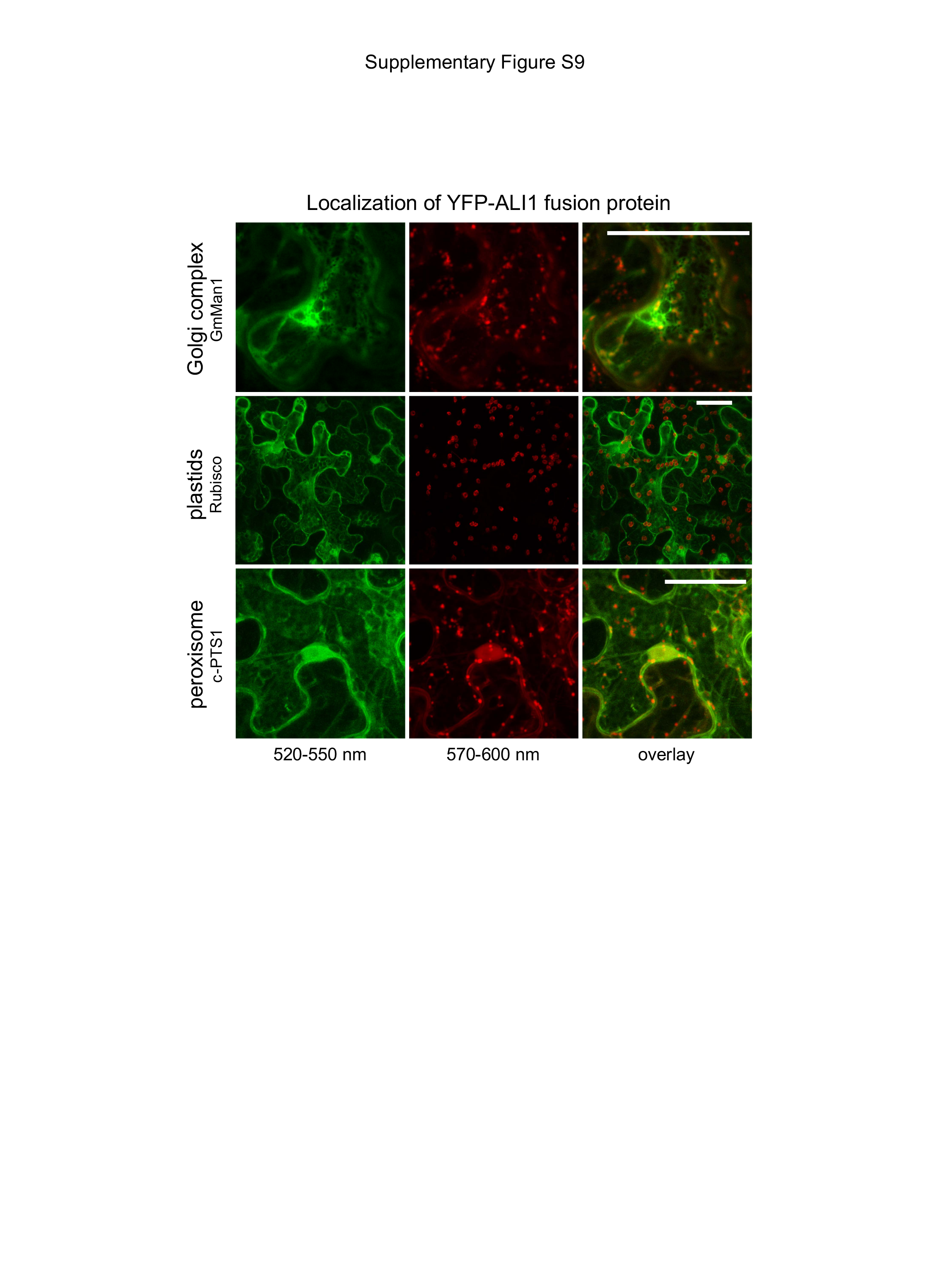

#### Supplemental Figure S4. ALI1 does not colocalize with Golgi, plastids and peroxisomes.

Plasmids carrying YFP-tagged ALI1 version and mCherry-marked proteins localizing to Golgi (GmMan1), plastids (RuBisCO) or peroxisome (PTS1) were co-transformed into *N. benthamiana* leaf epidermal cells by *Agrobacterium*-infiltration. The cells were analyzed 2 days after the infiltration using CLSM. The left panel shows in green, the fluorescence of the YFP-tagged ALI1, whereas the middle panel shows in red, the fluorescence of different mCherry-marked proteins colocalizing to different subcellular compartments (Supplemental Table S2). The right panel shows the corresponding merged image, where yellow color indicates a colocalization. Scale bar = 40 μm.

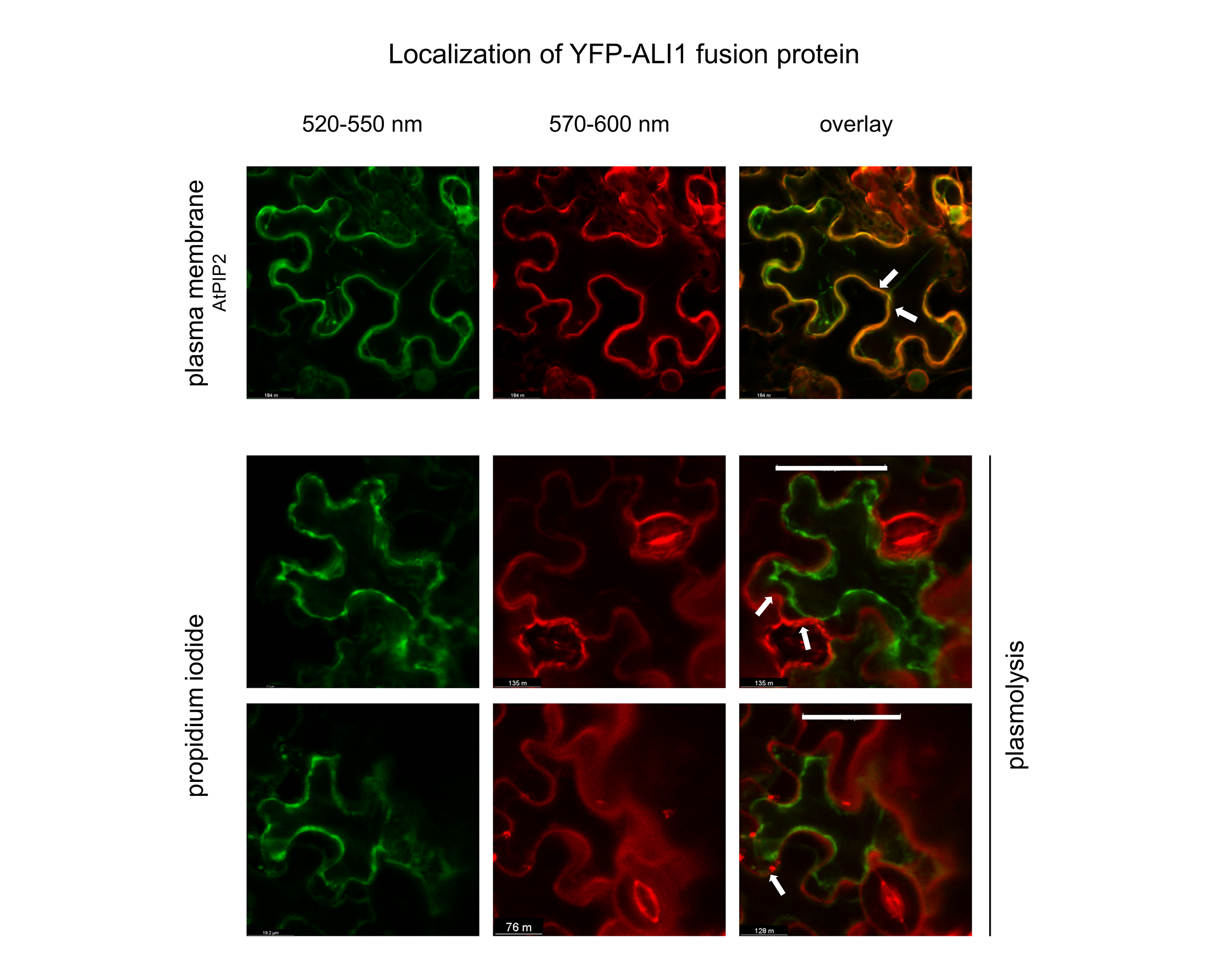

propidium iodide

#### Supplemental Figure S5. Localization of ALI1 upon plasmolysis.

Plasmids carrying YFP-tagged ALI1 version and mCherry-marked proteins localizing in PM (AtPIP2) were co-transformed to *N. benthamiana* leaf epidermal cells by *Agrobacterium*-infiltration. The cells were analyzed 2 days after the infiltration. The left panel shows in green the fluorescence of YFP-tagged ALI1 version, whereas the middle panel shows in red fluorescence of different mCherry-marked subcellular localizing proteins (upper panel), or the fluroscence of the propidium iodide. The right panel shows corresponding merged images where yellow color indicates colocalization. Scale bar = 40 µm. Arrows indicate the plasmolysis site (lower panels) or co-localization with AtPIP2 (upper panel).

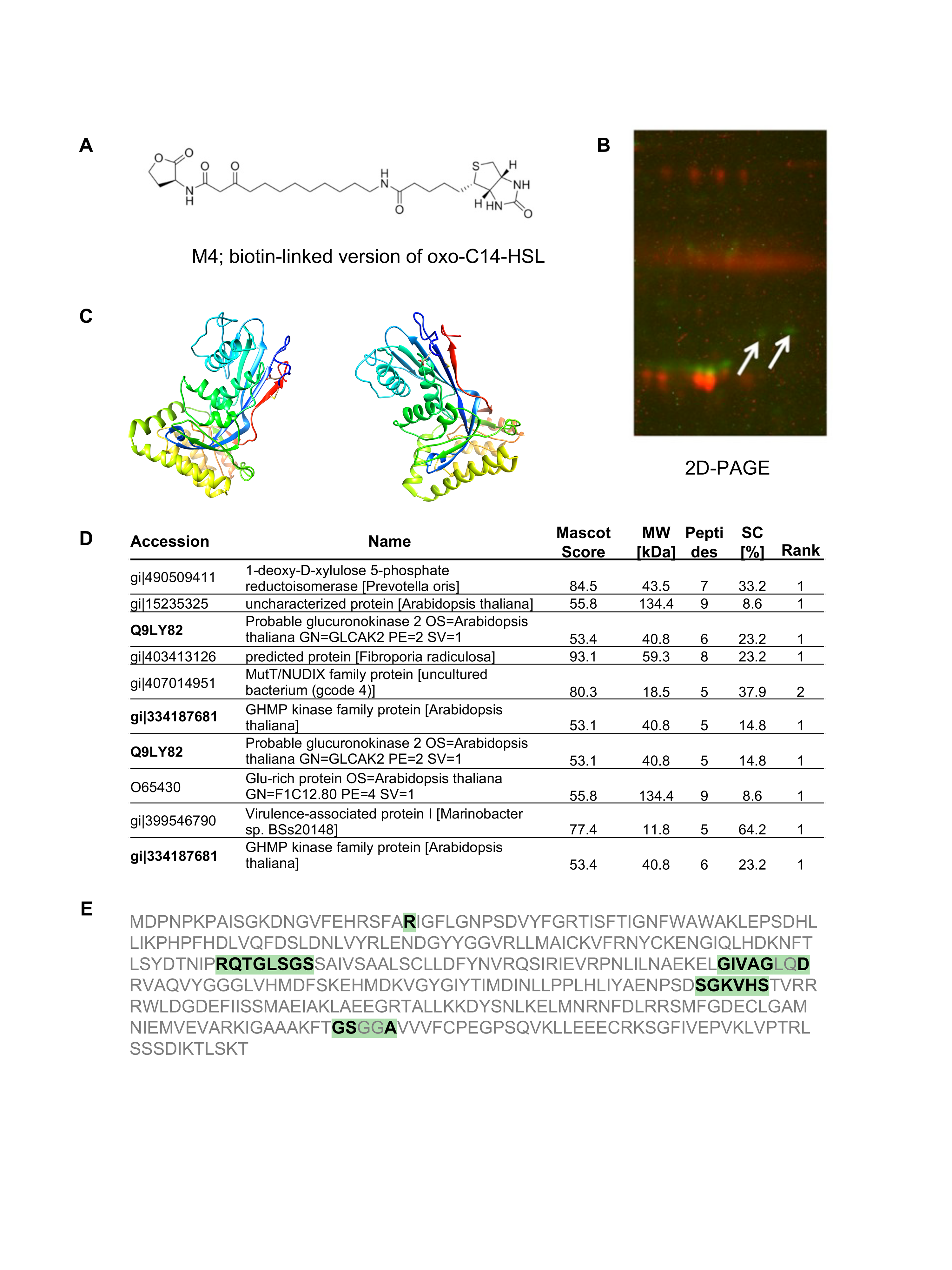

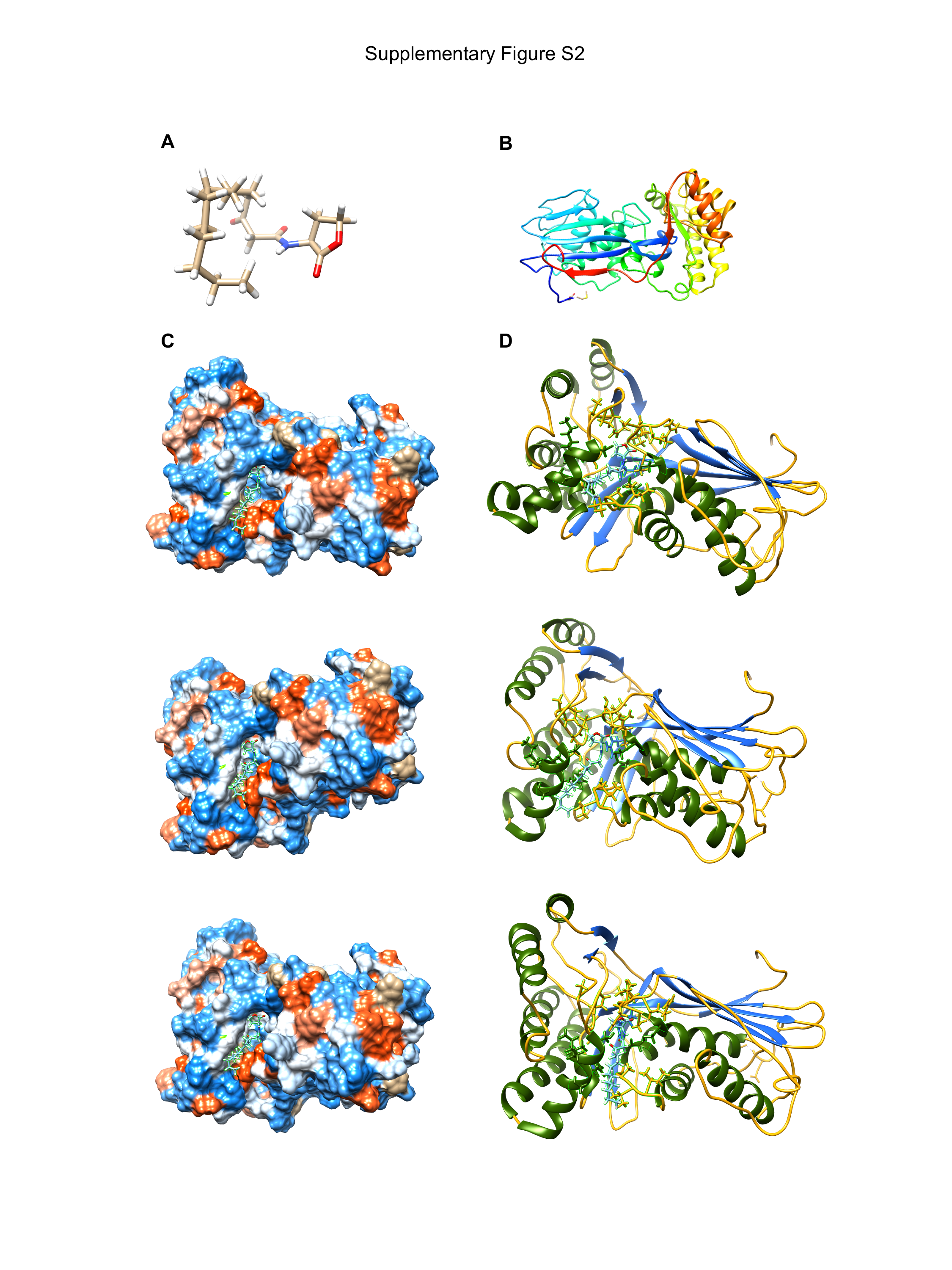

#### Supplemental Figure S6. Representative images of predicted docking between ALI1 protein and oxo-C14-HSL ligand.

The tertiary structure of (A) *N*-3-oxotetradecanoyl-*L*-homoserine lactone (oxo-C14-HSL) and (B) the predicted structure of ALI1 protein. The predicted hydrophobicity (C) and ribbon (D) structure of the ALI1 protein with the oxo-C14-HSL ligand from different angles. E, Amino acid sequences of ALI1, bold letters indicate residues with predicted distance of less than 5 Å from the ligand.

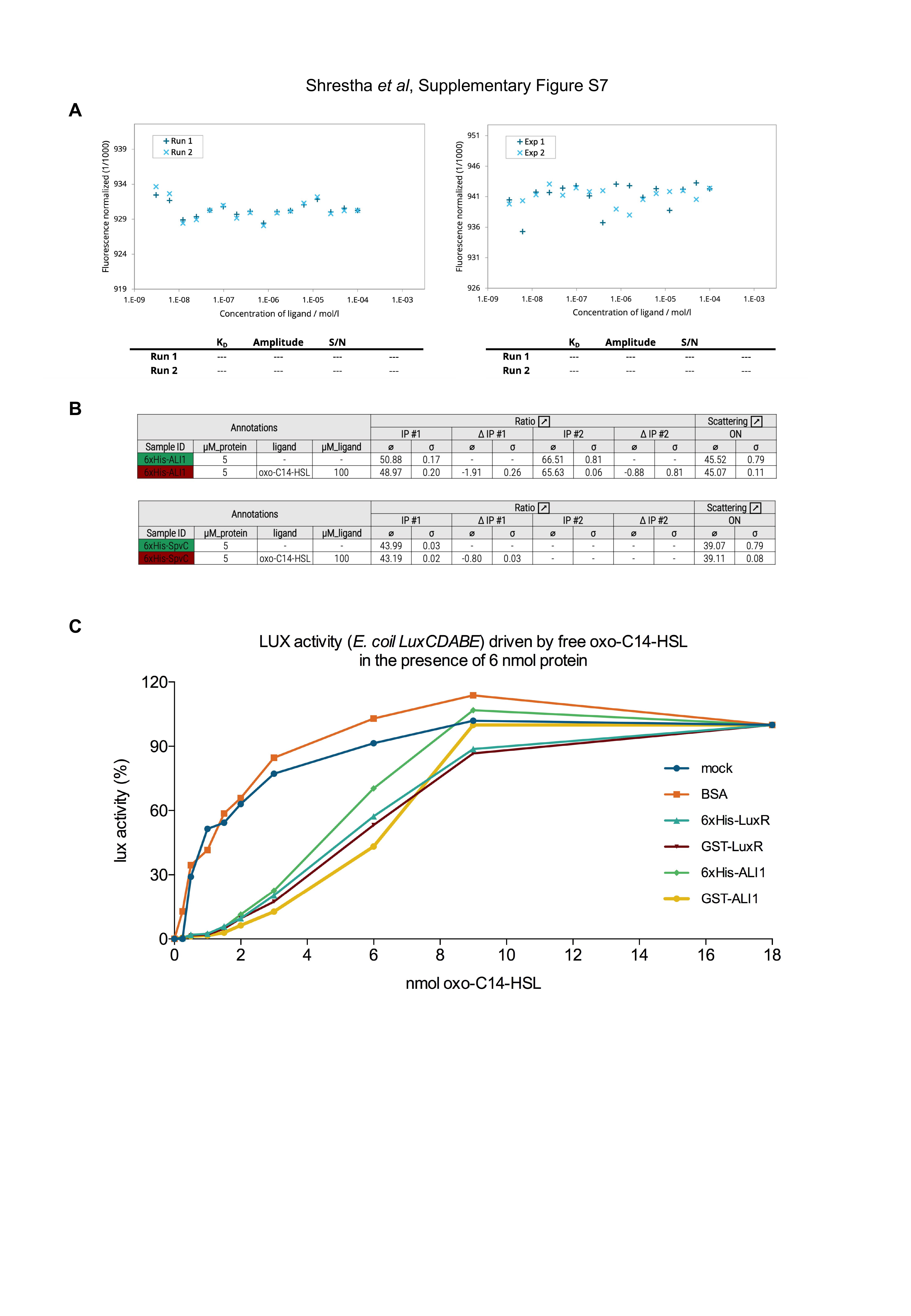

#### Supplemental Figure S7. Interaction between ALI1 and oxo-C14-HSL was missing in biophysical assays but indicated to be present in indirect binding assay.

A, MST binding assay missing to quantify the interaction study between fluorescently labeled proteins 6xHis-ALI1 (left) or 6xHis-SpvC (right) and ligand oxo-C14-HSL. The protein concentration was kept constant at 50 nM and the ligand oxo-C14-HSL was titrated from 100 µM to 3.05 nM. The difference in normalized fluorescence (%) was plotted for analysis of thermophoresis. Two experimental replicates were used for the analysis. B, Nano-DSF assay suggesting comparable thermal shifts in both proteins 6xHis-AI1 and 6xHis-SpvC in the presence of ligand oxo-C14-HSL which could possibly be an artifact. Both proteins and ligand were diluted to have a final concentration of 5 µM and 100 µM respectively. For both proteins, unfolding transition was assessed. Thermal shift was calculated by assessing difference of melting temperatures (Tm) between ligand-bound and apo-state of a protein. Two experimental replicates were used for the analysis. C, Luminescence activity assay indicating the binding of oxo-C14-HSL to ALI1. The binding capacity was assessed by determining the concentration of free oxo-C14-HSL using the *E. coil LuxCDABE* reporter strain after an overnight incubation with 6 nmol ALI1 or LuxR (positive control) and BSA (negative control) with different amounts of oxo-C14-HSL, as indicated.

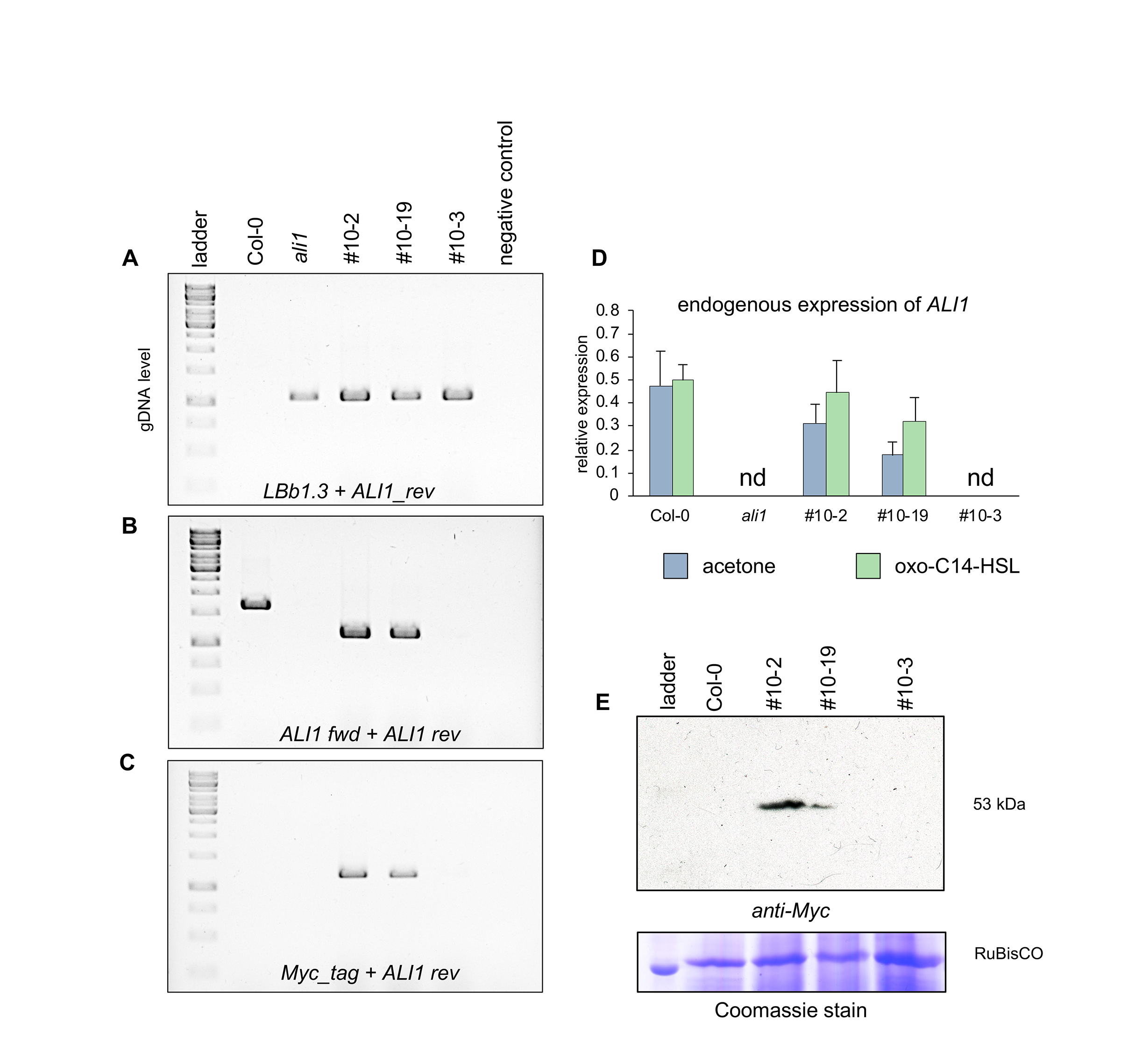

#### Supplemental Figure S8. The complemented lines of *ali1* mutant express ALI1.

Ethidium bromide stained gel indicating: (A) The presence of T-DNA insertion in *ALI1* gene of *ali1* mutant, two complemented lines of *ali1*: #10-2 and #10-19, and outcrossed line #10-3 but not in wild-type Col-0; (B) The presence of *ALI1* gene in Col-0 (genomic version) and two complemented lines of *ali1*: #10-2 and #10-19 (mRNA version); and (C) The presence of *ALI1* gene tagged with 10xMyc in two complemented lines of *ali1*: #10-2 and #10-19. Primers T-DNA left border specific LBb1-3 and ALI1 rev were used to detect the presence of T-DNA insertion in *ALI1* gene, gene-specific primers ALI1 fwd and ALI1 rev were used to detect the presence of *ALI1* gene and tag-specific primer Myc-tag and ALI1 rev were used to verify the presence of Myc-tagged *ALI1* gene. D, The endogenous expression level of *ALI1* in acetone and oxo-C14-HSL pretreated Col-0, *ali1*, two complemented lines of *ali1*: #10-2 and #10-19 and outcrossed line #10-3. Two-week old seedlings were transferred to ½-strength MS liquid medium supplemented with 6 µM oxo-C14-HSL or acetone control for three days. The abundance of *ALI1* gene transcript was normalized with *Ubiquitin ligase* (*At5g25760*) transcript. The bar represents mean and SD of three biological replicates. E, Western blot indicating the expression of Myc-tagged ALI1 in both complemented lines #10-2 and #10-19 but not in outcross line #10-3. Representative blot is shown.

### Supplemental Tables

#### Supplemental Table S1. Primers used in the study

| **Oligonucleotide** | **Sequence (5´–3´)** | **Reference** |
| --- | --- | --- |
| ALI1 fwd | GGA GAT AGA ACC ATG GAT CCG AAT CCT AAA CCG | This study |
| ALI1 rev | TCC ACC TCC GGA TCM TGT TTT TGA TAA TGT CTT AAT ATC AGA AC | This study |
| U5 fwd | GGG GAC AAG TTT GTA CAA AAA AGC AGG CTT CGA AGG AGA TAG AAC CAT G | ATOME Project |
| U3 rev | AGA TTG GGG ACC ACT TTG TAC AAG AAA GCT GGG TCT CCA CCT CCG GAT C | ATOME Project |
| DNR5 | CTG GCA GTT CCC TAC TCT CG | (Hilson et al., 2004) |
| DNR3 | GAT GGT CGG AAG AGG CAT AA | (Hilson et al., 2004) |
| GTW1 | TAG CTT CCT TAG CTC CTG AAA ATC TCG | ATOME Project |
| GTW2 | GGG AAT ATA AAT GTC AGG CTC CCT TA | ATOME Project |
| 35S-promoter | TTC GCA AGA CCC TTC CTC TAT A | universal |
| Actin fwd | GGT CGT ACA ACC GGT ATT GTG CTG G | C. Forzani (*personal comm.)* |
| Actin rev | TTG GAG ATC CAC ATC TGC TGG AAT G | C. Forzani (*personal comm.)* |
| T7 | TAA TAC GAC TCA CTA TAG GG | universal |
| UBQ fwd | GCT TGG AGT CCT GCT TGG ACG | (Schenk et al., 2014) |
| UBQ rev | CGC AGT TAA GAG GAC TGT CCG GC |  |
| WRK22 fwd | ATC TCC GAC GAC CAC TAT TG | (Schenk et al., 2014) |
| WRK22 rev | TCA TCG CTA ACC ACC GTA TC |  |
| WRK29 fwd | TCC GGT ACG TTT TCA CCT TC | (Schenk et al., 2014) |
| WRK29 rev | AGA GAC CGA GCT TGT GAG GA |  |
| GST6 fwd | GCA TGT TCG GCA TGA CCA CTG | (Schenk et al., 2014) |
| GST6 rev | GCA CCT TGG AGT CAG TAC CC |  |
| Hsp70 fwd | CGC CAA CGA TCA AGG CAA CC | (Schenk et al., 2014) |
| Hsp70 rev | GCT TCT CAC CTG GAC CGG AA |  |
| ALI1 ins1 | GAT CTT ACG TGC CAC TTC CAC | This study |
| ALI1 ins2 | TTT TGA TTG AAA GAT TTG TGG C | This study |
| ALI1 qPCR fwd | GAC GGG GCT TTC AGG TTC TA | This study |
| ALI1 qPCR rev | AGA CTT GAG CAA CAC GGT CT | This study |
| LBb1-3 | ATT TTG CCG ATT TCG GAA C | Salk Institute |
| Myc-tag | AAT CTC CGA GGA AGA CTT GAA C | This study |

#### Supplemental Table S2. Fluorescence tagged strains used in the localization study

Localization markers used in this study in the *Agrobacterium*-mediated transformation.

| **Strain** | **Remarks** | **Localization** | **Reference** |
| --- | --- | --- | --- |
| *A. tumefaciens* (LBA4404) | pBin20 containing mCherry with the ATWAK2 signal peptide and HDEL-motif | endoplasmic reticulum | (Nelson et al., 2007) |
| *A. tumefaciens* (LBA4404) | pBin20 containing mCherry fused to full-length coding region of AtPIP2A, a plasma membrane aquaporin (Cutler et al., 2000) | plasma membrane | (Nelson et al., 2007) |
| *A. tumefaciens* (LBA4404) | pBin20 containing mCherry fused to the C-terminus of c-TIP, an aquaporin of the vacuolar membrane (Saito et al., 2002) | tonoplast | (Nelson et al., 2007) |
| *A. tumefaciens* (LBA4404) | pBin20 containing mCherry fused to peroxisomal targeting signal1 (PTS1, Ser-Lys-Leu) at its C-terminus | peroxisome | (Nelson et al., 2007) |
| *A. tumefaciens* (LBA4404) | pBin20 containing mCherry fused to first 29 aa of yeast (*Saccharomyces cerevisiae*) cytochrome c oxidase IV | mitochondria | (Nelson et al., 2007) |
| *A. tumefaciens* (LBA4404) | pBin20 containing mCherry fused to targeting sequence (first 79 aa) of the small subunit of tobacco rubisco | plastids | (Nelson et al., 2007) |
| *A. tumefaciens* (LBA4404) | pBin20 containing mCherry fused to cytoplasmic tail and transmembrane domain (first 49 aa) of GmMan1, soybean a-1,2-mannosidase I | Golgi | (Nelson et al., 2007) |

### Supplemental Datasets

**Supplemental** Dataset S1. Genes differentially expressed (upregulated) in Col-0 after 3-day pretreatment with oxo-C14-HSL and additional challenge with 100 nM flg22 for 2 h

**Supplemental** Dataset S2. Genes differentially expressed (downregulated) in Col-0 after 3-day pretreatment with oxo-C14-HSL and additional challenge with 100 nM flg22 for 2 h

**Supplemental** Dataset S3. Genes differentially expressed (upregulated) in *ali1* after 3-day pretreatment with oxo-C14-HSL and additional challenge with 100 nM flg22 for 2 h

**Supplemental** Dataset S4. Genes differentially expressed (downregulated) in *ali1* after 3-day pretreatment with oxo-C14-HSL and additional challenge with 100 nM flg22 for 2 h
